# Engineering gene expression dynamics via self-amplifying RNA with drug-responsive non-structural proteins

**DOI:** 10.1101/2025.06.08.658495

**Authors:** Parisa Yousefpour, Justin R. Gregory, Kristen Si, Jan Lonzaric, Yingzhong Li, Junmin Wang, Kashif Qureshi, Amir Ledbetter, Ashley A. Lemnios, Jonathan Dye, Tanaka K. Remba, Rachel Yeung, Melissa Güereca, Linette Rodriguez, Yuebao Zhang, Shengwei Wu, Yizhou Dong, Ron Weiss, Darrell J. Irvine

## Abstract

The design of gene therapies with drug-regulatable expression of therapeutic payloads is of interest for diverse applications. We hypothesized that a regulated expression system based on alphavirus-derived self-amplifying RNAs (saRNAs), which encode 4 non-structural proteins (nsPs) that copy the RNA backbone to enable sustained expression, would have advantages in safety and simplicity of delivery. Here we designed saRNAs where payload expression is regulated by the FDA-approved drug trimethoprim (TMP), by fusing TMP-responsive degradation domains (DDs) to nsPs to regulate RNA self-amplification. Screening a library of nsP-DD fusions, we identified an optimal design with DDs fused to nsP2, nsP3, and the payload, achieving a high fold-change in expression level in response to TMP and low expression in the off state. In mice, this saRNA circuit enabled diverse dynamic expression patterns in response to oral TMP. Implementing this circuit for controlled expression of an HIV antigen, an escalating TMP regimen significantly enhanced germinal center responses critical for B cell affinity maturation. This drug-regulated RNA technology holds potential for vaccines, immunotherapies, and gene therapies.

---

Gene therapies have made steady progress in the clinic, but genetic systems where expression of a therapeutic payload can be modulated over time are of great interest. For example, in cancer immunotherapy, the order and timing of immunostimulatory cues can significantly impact treatment efficacy and toxicity.^1–4^ Similarly, in vaccine development, changes in antigen availability over time can modulate the immune response, with escalating antigen doses over a few weeks leading to stronger humoral and T cell responses.^5–7^ In recent years, several approaches for regulated gene expression have advanced to human testing, each offering unique advantages but also facing significant limitations. Viral vector systems, such as adeno-associated viruses (AAVs) or lentiviruses, are widely used,^8^ but face anti-vector immunity, inflammatory toxicities,^9–12^ and concerns about insertional mutagenesis.^13,14^ Tetracycline-inducible systems offer reversible control in response to small-molecule drugs but often exhibit leaky expression. Other systems that have shown promise include tetracycline-inducible systems, which offer reversible control of gene expression in response to small-molecule drugs, but often suffer from leaky expression.^15–17^ CRISPR-based approaches, including CRISPRa and CRISPRi, allow for controlling gene expression but face delivery challenges and potential off-target effects.^18,19^ Rapamycin-inducible and progesterone analog-inducible systems provide rapid, dose-dependent control but may have unintended effects on endogenous signaling pathways.^20,21^

A regulated RNA-based gene expression system offers theoretical advantages over these existing approaches, including (1) no concern of chromosomal integration, (2) simplified delivery using clinically validated delivery systems such as lipid nanoparticles (LNPs),^22^ and (3) lack of vector-related toxicity/immunogenicity issues. Current mRNA designs, even employing modified bases for enhanced stability and reduced innate immune recognition, are not well-suited for this purpose as they have relatively short-lived expression *in vivo.* However, in recent years, self-amplifying RNAs (saRNAs or replicons) derived from viral genomes have emerged as an alternative for long-lived therapeutic gene expression. Capable of autonomous replication within host cells, saRNAs can achieve sustained protein expression lasting up to many weeks from low doses of RNA, making them an attractive platform techonlogy.^23–29^ saRNAs are engineered from positive-strand RNA viruses, like alphaviruses and flaviviruses, by replacing the structural protein genes with a gene of interest (GOI) while retaining the non-structural proteins (nsPs) responsible for RNA replication.^30^ This design enables high-level, prolonged expression of the desired protein without producing infectious viral particles. saRNAs are in clinical testing for therapeutic vaccines against cancer^31–33^ and solid tumor immunotherapy,^34^ and saRNA-based COVID-19 vaccines have received clinical approval in several countries.^35–37^ Replicons that alter payload expression in response to a small-molecule drug via a bacterial degradation domain have been reported, where payload expression is high in the absence and reduced in the presence of the drug.^38^ However, this approach provides very leaky expression of the reporter gene in the “off” state, and a drug-induced “on” state would be more attractive in many applications.

We hypothesized that regulation of not only the payload gene but also the self-amplifying mechanism of replicon RNA itself would enable more robust control over expression. To achieve this, here we developed an approach for drug-regulated saRNAs by fusing a degradation domain (DD) responsive to the FDA-approved small molecule antibiotic trimethoprim (TMP)^39,40^ directly with the nsPs forming the replicase complex, enabling TMP-mediated control of replicon copy number within the cell. We found that saRNAs carrying DD-modified nsPs retained their self-amplifying capacity and robust, long-lived gene expression. An optimal regulated saRNA employing DD-fused nsP2 and nsP3 proteins together with a DD-regulated payload achieved high-fidelity, reversible TMP-regulated expression *in vivo*, allowing modulation of payload expression over several weeks. As a proof-of-concept application example, we show how this engineered gene circuit could be used to achieve temporally programmed escalating expression of a model HIV antigen, leading to augmented expansion of germinal center B cells. By combining sustained expression capabilities of saRNAs with tunable replication control, we arrive at a platform enabling high-fidelity drug-regulated payload expression with low leakiness, a high-level “on” state, and reversible long-term expression control.

## Results

### Fusion of DD to nsPs effectively controls transgene expression from saRNAs

We used a Venezuelan equine encephalitis virus (VEEV)-derived saRNA with the structural proteins encoded under the virus’s subgenomic promoter replaced by a payload GOI as the backbone for our regulated expression platform. VEEV saRNA delivered into the cytoplasm of host cells rapidly self-amplifies via four nsPs, which generate negative and positive-strand copies of the entire replicon and transcribe the subgenome encoding the GOI (**Fig. 1a**). We hypothesized that fusing nsPs with a destabilization domain (DD) responsive to the small molecule antibiotic trimethoprim (TMP)^40^ could regulate RNA replication, achieving high-fidelity on/off control and a wide dynamic range of payload expression.

**Figure 1:**
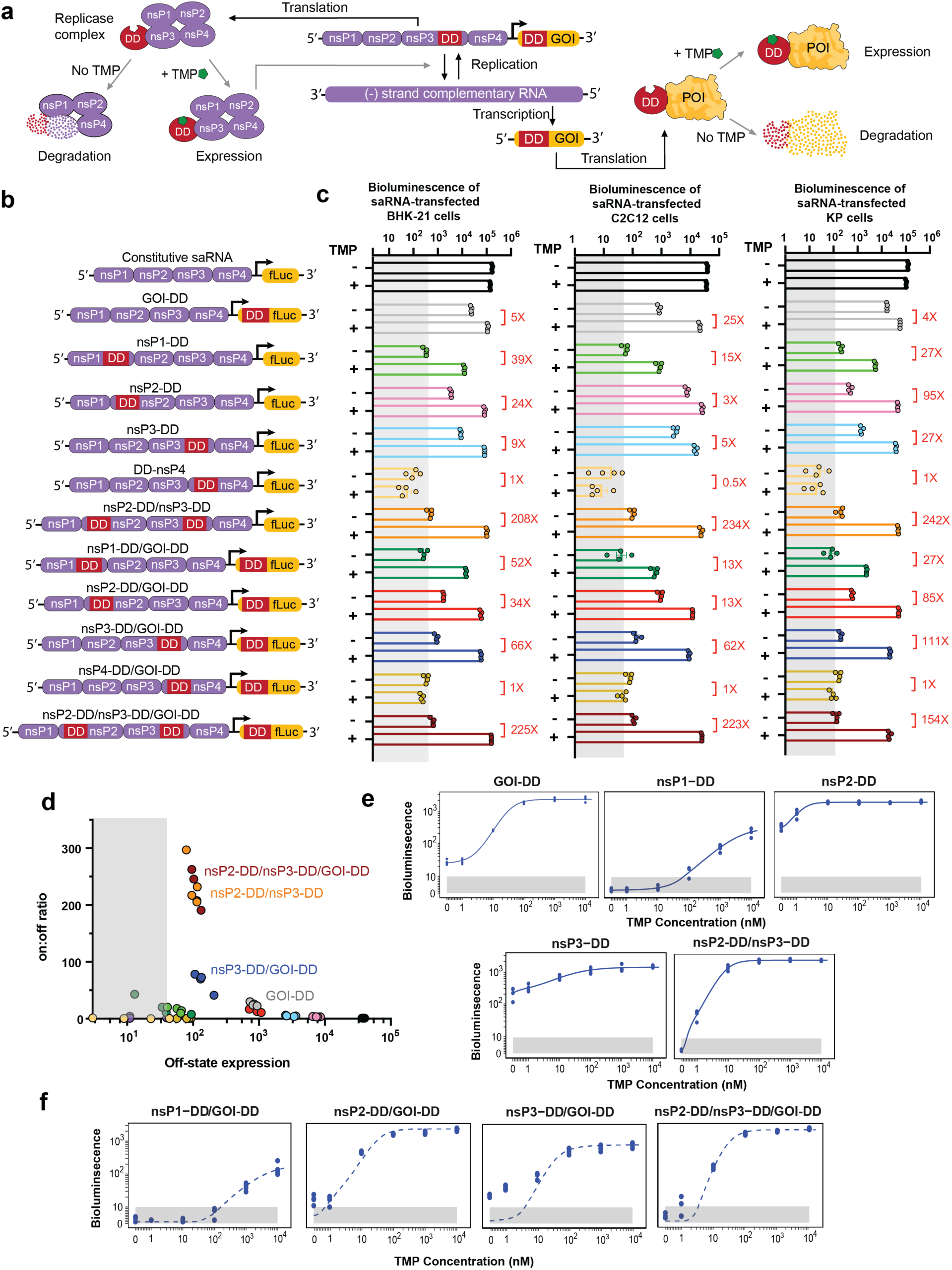
Regulation of transgene expression from saRNAs by small molecule drug trimethoprim (TMP). a, TMP-regulated saRNAs are created by fusing a TMP-responsive degradation domain (DD) to the non-structural proteins (nsPs) of the saRNA backbone and/or the gene of interest (GOI). Fusion to nsPs enables regulation of saRNA and subgenome RNA copy numbers, while fusion to the GOI controls protein stability post-translation. POI, protein of interest. **b,** Schematics of gene circuits created by fusing DD to each of the four nsPs forming the replicase complex, with or without an additional DD fused to the GOI. Two additional circuits were created by fusing DD to both nsP2 and nsP3, with or without an additional DD fused to the GOI. **c,** Regulation of transgene expression from different DD-based saRNA circuits in response to TMP. C2C12 mouse myocytes, BHK-21, and KP lung cancer cells were transfected with different saRNA gene circuits encoding firefly luciferase (fLuc) as a reporter cargo and were cultured with or without TMP post-transfection. FLuc expression was assessed by bioluminescence at 24 hours post-transfection. The shaded gray area indicates background bioluminescence from control cells electroporated without replicon. **d,** Scatter plot of on:off ratio versus off-state bioluminescence signal in C2C12 cells transfected with genetic circuits, measured at 24 hours post-transfection. Circuits in the upper-left quadrant, exhibiting high on:off ratios and low “off” state expression are considered optimal for precise control of gene expression. Shaded area indicates background bioluminescence levels. Colors correspond to those in panels b-c. Data are presented as mean ± SEM with indicated *n*. **e,f,** Fitted/predicted dose-response curves and experimental data for DD-based circuits. C2C12 cells were transfected with saRNA gene circuits and incubated with different concentrations of TMP. FLuc expression was assessed by bioluminescence at 24 hours post-transfection. Solid curves represent the fitted models, dashed curves represent the predictions, and dots correspond to the experimental data. The y-axis displays fLuc intensity on a semi-log scale, with a linear scale from 1 to 10 (shaded in gray to indicate the background fLuc intensity) and a log scale for values above the background intensity of 10. The x-axis displays TMP concentration on a semi-log scale, with a linear scale from 0 to 1 and a log scale for values above 1. **(e)** Curves were fitted for fLuc-DD, nsP1-DD/fLuc, nsP2-DD/fLuc, nsP3- DD/fLuc, and nsP2-DD/nsP3-DD/fLuc circuits using a 4PL model. **(f)** Prediction curves were generated for nsP1- DD/fLuc-DD, nsP2-DD/fLuc-DD, nsP3-DD/fLuc-DD, and nsP2-DD/nsP3-DD/fLuc-DD circuits based on the assumption of modularity.

To test this idea, we created a library of mutant saRNAs, fusing a mutant dihydrofolate reductase (DHFR) DD to one or more of the four nsPs that constitute the saRNA replicase complex (**Fig. 1a-b**). This DD promotes degradation of its fusion partner unless stabilized by TMP. We predicted that DD fusion would disrupt the replicase complex and suppress saRNA replication unless TMP was present to stabilize the fusion (**Fig. 1a**). Fusion at the N-terminus or C-terminus of each nsP was strategically selected to minimize disruption of conserved backbone sequences and essential functional domains:^41–45^ for nsP1, DD was fused to the N-terminus to maintain the integrity of conserved sequence elements (CSEs) at the 5’ end of its RNA sequence;^46^ in nsP2, DD was also fused to the N-terminus to prevent interference with its C-terminal protease activity and substrate recognition functions; for nsP3, DD was fused to the C-terminus, as this region is a disordered, non-conserved domain.^47^ Additionally, the cleavage site between nsP2 and nsP3 is tightly regulated.^44,48,49^ Positioning DD at the N-terminus of nsP2 and C-terminus of nsP3, away from this site, minimizes potential interference with replicase processing and function. For nsP4, DD was fused to the N-terminus, preserving the C-terminal CSE and subgenomic promoter sequence.^46^

In a second set of designs, an additional DD was fused directly to the GOI encoded by the saRNA subgenome **(Fig. 1a-b)**. This library of engineered saRNAs were prepared by *in vitro* transcription at high purity (**Supplementary Fig. S1**).

To test the expression behavior of these constructs, we first assessed the behavior of saRNAs encoding firefly luciferase (fLuc) as a reporter GOI *in vitro* in BHK-21 cells that are highly permissive for saRNA expression.^24,25^ fLuc expression was measured 24 hours post-transfection by luminescence assays with or without 10 µM TMP. Fusing DD to the payload gene alone enabled strong “on” (+TMP) state expression nearing the level observed for the constitutive saRNA with no DD introduction, but was very leaky in the “off” (-TMP) state, with expression well above the background level of non-expressing cells, and only a 4-fold change from off-to-on state expression levels (**Fig. 1c**). Examining first constructs bearing a DD fused to a single nsP, we found that nsP1-DD exhibited TMP-sensitive expression with a better “off” state than the fLuc-DD control but did not reach the levels of “on” state expression achieved by fLuc-DD or the constitutive control saRNA. DD fused to nsP2 or nsP3 achieved high on-state expression but were also highly leaky in the “off” state, while nsP4-DD fusions did not express at all. Construction of a control saRNA using wild type-DHFR, which differs from DD by only two mutations, led to loss of TMP regulation, as expected (**Extended Data** Fig. 1). We next tested tandem DD constructs with DD fused to one nsP and the payload gene. Double-DD nsP-DD/GOI-DD saRNAs where DD was fused to nsP1, nsP2, or nsP4 showed similar behavior as the single nsP-DD circuit designs. However, nsP3-DD/fLuc-DD exhibited enhanced regulation over either of the single DD constructs, with high “on” state expression and lower “off” state background, providing an ∼70-fold dynamic range in on/off state expression. In an attempt to further increase the expression dynamic range, we finally evaluated saRNAs where DD was fused to both nsP2 and nsP3. This tandem nsP-DD construct had low off-state background expression and achieved an impressive 210-fold change in fLuc signal in the “off” vs. “on” state, further slightly enhanced by additionally fusing the payload gene with DD **(Fig. 1c)**. We also tested expression in murine C2C12 myoblast cells (representing a tissue target, muscle, of interest for gene and vaccine delivery), a murine mutant-Kras lung cancer cell line,^50^, and human HEK293T cells. These cells showed similar patterns of regulated saRNA expression (**Fig. 1c** and **Extended Data** Fig. 2). The nsP2-DD/nsP3-DD/GOI- DD, nsP2-DD/nsP3-DD, and nsP3-DD/GOI-DD circuits consistently demonstrated the most optimal control of transgene expression, characterized by low off-state expression and high on-to-off expression ratios **(Fig. 1d)**.

To further quantitatively characterize the constructs, we conducted dose titration experiments measuring fLuc expression vs TMP concentration in C2C12 cells. We fit the data from the single-nsP-DD and GOI-DD fusion constructs to four-parameter logistic (4PL) curves (see Methods). The 4PL model was selected for its ability to capture the sigmoidal relationship inherent in the system, enabling precise parameterization of minimum and maximum expression levels, inflection point, and slope (**Supplementary Table S1**). The models fit the data well (**Fig. 1e**), yielding R^2^ values of 0.996, 0.971, 0.960, 0.915, and 0.996 for GOI-DD, nsP1-DD, nsP2-DD, nsP3-DD, and nsP2-DD/nsP3-DD circuits, respectively. Next, we assumed a multiplicative effect for circuits combining GOI- DD with nsP-DD designs, and examined the predictive power of the model by comparing computational predictions with experimental fLuc dose-response data.^51–53^ Three of the four resulting predictions showed good agreement with experimental observations (**Fig. 1f**), with R^2^ values of 0.843, 0.922, and 0.843 for nsP1-DD/GOI-DD, nsP2- DD/GOI-DD, and nsP2-DD/nsP3-DD/GOI-DD circuits, respectively. For the nsP3-DD/GOI-DD circuit, the predictions underestimated the basal expression level. The overall agreement between predicted and observed results supports a multiplicative effect between the GOI-DD and most nsP-DD components, suggesting predictable outputs for more complex designs. For the nsP3-DD/GOI DD circuit, the observed deviation suggests additional factors at play that would require further exploration to fully understand.

We next assessed the on/off behavior in C2C12 cells for engineered saRNAs expressing mCherry as the GOI, where relative quantities of expressed payload protein are directly proportional to fluorescence signal and expression analysis at the single-cell level is straightforward. Consistent with findings from the fLuc-encoding saRNAs, fusion of DDs to nsP1, nsP2, and nsP3 enabled strong regulatory control, with significant differences in the percentages of mCherry^+^ cells and levels of mCherry expression between the "on" and "off" states (**Fig. 2a**, **Extended Data** Fig. 3). Circuits combining regulated nsP3 with either a regulated payload or regulated nsP2 achieved the highest on/off expression ratios and lowest background in the absence of TMP **(Fig. 2b)**. Notably, unlike the fLuc reporter, these circuits had essentially no detectable background expression of mCherry in their “off” states.

**Figure 2:**
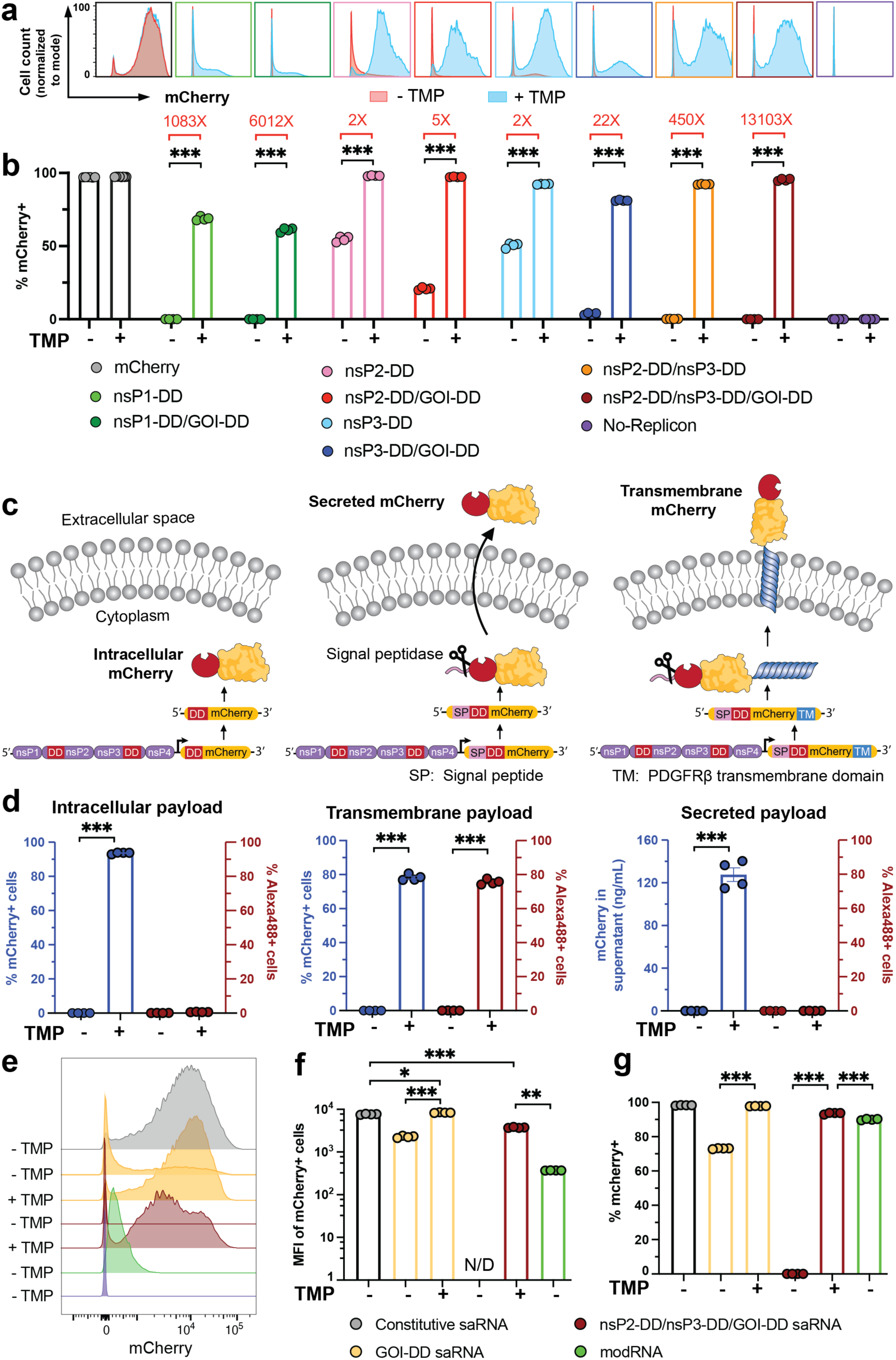
Versatility of gene circuits. a,b, Expression for fluorescent reporter mCherry from saRNA gene circuits. C2C12 cells were transfected with saRNA gene circuits encoding mCherry. At 24 hours post-transfection, and gene expression was evaluated at the individual cell level by flow cytometry. Shown are the representative flow cytometry histograms **(a)**, and percentage of cells expressing mCherry **(b)**. **c,d,** Effectiveness of nsP2-DD/nsP3-DD/GOI-DD circuit for different protein formats: C2C12 cells were transfected with nsP2/nsP3- DD/GOI-DD saRNA circuit encoding mCherry in the transmembrane, secreted, and intracellular formats **(c)**. At 24 hours post-transfection, mCherry expression was assessed by ELISA for the secreted format and by flow cytometry for the intracellular and transmembrane formats. For the transmembrane format, cells were stained with an Alexa488-conjugated anti-mCherry antibody to distinguish mCherry on the cell surface from that inside the cells **(d)**. **e,f,g,** Comparison of mCherry expression levels. Shown are the representative flow cytometry histograms **(e)**, percentages of mCherry+ cells **(f)**, and mean fluorescent intensity (MFI) of mCherry+ cells **(g)** in C2C12 cells transfected with mcherry-encoding nsP2-DD/nsP3-DD/GOI-DD saRNA, constitutive saRNA, and modRNA. N/D: Not Detected, indicating no or too few mCherry+ cells to reliably calculate MFI. Data are presented as mean ± SEM with indicated *n*. Statistical comparisons were performed using one-way ANOVA, followed by Tukey’s post-hoc test. *, P < 0.05; **, P < 0.01; ***, P < 0.001.

Interestingly, TMP-dependent regulation was maintained even when gene circuit-incorporating saRNAs were co-transfected with wildtype saRNAs that express the replicase independently of TMP. In C2C12 cells co-electroporated with constitutively-expressing GFP and TMP-regulated mCherry-expressing saRNAs containing various nsP-DD gene circuits, mCherry expression remained under TMP control, even in the absence of a DD directly fused to mCherry. However, the degree of regulation was diminished, with a lower fold change in the percentage of mCherry^+^ cells between the "on" and "off" states compared to cells transfected exclusively with the regulated saRNA **(Extended Data** Fig. 4).

To explore diverse payload types, we created nsP2-DD/nsP3-DD/GOI-DD saRNAs encoding mCherry as an intracellular, secreted, or transmembrane (TM) payload (**Fig. 2c**). Surface-expressed mCherry-DD was detected by flow cytometry using Alexa488-conjugated anti-mCherry antibodies, while secreted mCherry was quantified by ELISA. All expression forms showed essentially zero background expression without TMP, and robust expression with TMP (**Fig. 2d**). For intracellular mCherry, we compared the percentage and median fluorescent intensity (MFI) of mCherry^+^ cells transfected with the nsP2-DD/nsP3-DD/GOI-DD circuit to those transfected with a regulated circuit with DD only fused to GOI, non-regulated saRNA, and non-replicating base-modified mRNA. The MFI of mCherry from the regulated saRNA circuit was only 2-fold lower than that of a non-engineered, constitutive saRNA, yet it was 10-fold higher than non-replicating mRNA, demonstrating the expression advantage of saRNA **(Fig. 2e- f)**. Furthermore, the nsP2-DD/nsP3-DD/GOI-DD circuit exhibited tight regulation of mCherry expression with negligible mCherry^+^ cells in the “off” state, whereas the GOI-DD circuit displayed considerable leakiness, with ∼70% of cells expressing mCherry in the “off” state **(Fig. 2g)**. Altogether, these experiments identified nsP3- DD/GOI-DD and nsP2-DD/nsP3-DD saRNAs as interesting candidate circuits for further study.

### Regulated nsPs enable control of saRNA backbone and subgenomic RNA levels by TMP

To determine how TMP regulation affected RNA transcript levels, we performed quantitative PCR (qPCR) to quantify RNA copy numbers for the saRNA backbone and subgenomic RNA (encoding the fLuc payload) for the best-performing circuits. nsP3-DD and nsP3-DD/fLuc-DD saRNAs showed 2 orders of magnitude lower backbone and payload RNA copies in the “off” state compared to the non-regulated saRNA, while levels in the + “on” state were comparable (**Fig. 3a**). By contrast, nsP2-DD/nsP3-DD saRNAs showed even lower backbone and subgenomic RNA levels in the “off state, compared to the non-regulated saRNA. Unexpectedly, in the presence of TMP, both backbone and subgenomic RNA levels were over two orders of magnitude higher for nsP2-DD/nsP3- DD saRNAs compared to non-regulated constructs (**Fig. 3b**), suggesting enhanced polymerase activity with DD fusion to both nsP2 and nsP3.

**Figure 3:**
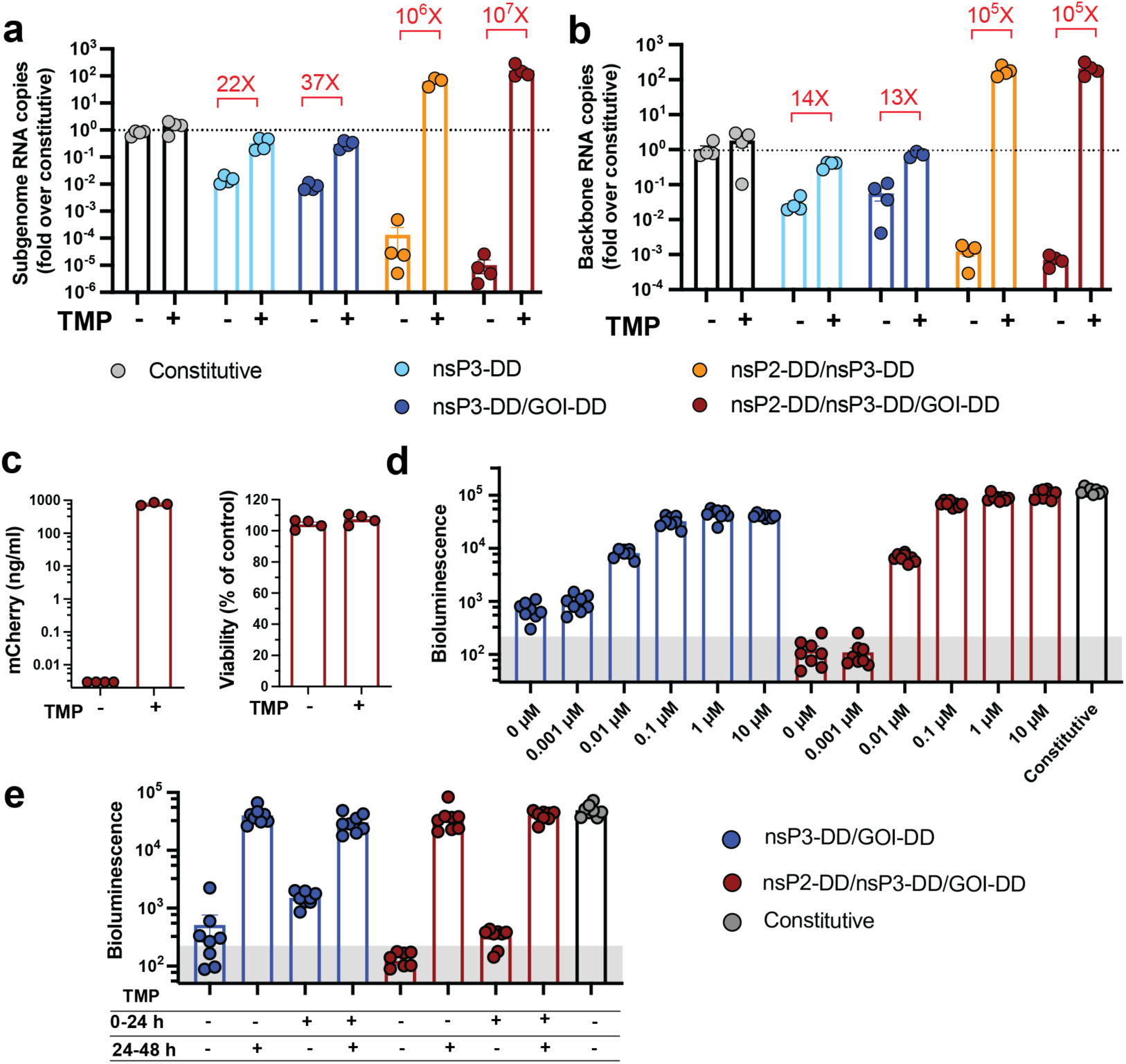
Quantitative and functional analysis of gene circuit dynamics in vitro. a,b, Relative quantification of saRNA and subgenomic RNA copy numbers. RNAs were extracted from C2C12 cells transfected with saRNAs, and following cDNA synthesis, qPCR was performed to quantify relative RNA levels for **(a)** saRNA (using primers targeting the nsP3 gene, and **(b)** subgenomic RNA. **c,** Toxicity and expression control of nsP2-DD/nsP3-DD/GOI- DD circuit in differentiated myofibers in vitro. Differentiated myofibers were treated with LNPs encapsulating mCherry-encoding circuits. Cell viability and mCherry expression were assessed at 24 hours post transfection by CellTiter glow and mCherry ELISA assays, respectively. **d,** Tunability of nsP3-DD/GOI-DD and nsP2-DD/nsP3- DD/GOI-DD circuit. Adjusting Transgene Expression Level by fine-tuning TMP concentration: fLuc expression from the nsP3-DD/GOI-DD and nsP3-DD/GOI-DD circuit was assessed by culturing differentiated myofibers with different TMP concentrations after electroporation and measuring bioluminescence after 24 hours. **e,** Reversibility of nsP3-DD/GOI-DD and nsP2-DD/nsP3-DD/GOI-DD circuits: Differentiated myofibers transfected with the saRNAs were incubated with or without TMP for 24 hours. Subsequently, cells initially treated with 0 or 10 μM TMP were switched to 10 or 0 μM TMP, respectively. FLuc expression was measured by bioluminescence at 48 hours post-transfection (24 hours after switching). The shaded gray area indicates background bioluminescence from control untreated cells. Data are presented as mean ± SEM with indicated *n*.

### saRNAs with regulated nsPs expression optimize gene expression control in muscle fibers

We next tested the behavior of the optimal regulated circuits in C2C12 cells differentiated from myoblasts into muscle fibers,^54,55^ an important transfection target for vaccines and gene therapies.^56–59^ C2C12 myofibers were transduced with saRNAs containing the high-fidelity nsP2-DD/nsP3-DD/mCherry-DD or nsP3-DD/GOI-DD constructs using LNPs. Importantly, these circuits showed similarly effective expression regulation in response to TMP in myofibers, with minimal toxicity and nearly 5 orders of magnitude difference in mCherry levels between the “on” and “off” states **(Fig. 3c)**. Titration of TMP added to myofiber cultures showed dose-dependent induction of payload expression in response to the drug for both nsP2-DD/nsP3-DD/GOI-DD and nsP3-DD/GOI-DD circuits, with expression plateauing at ∼0.1 µM TMP (**Fig. 3d**).

To assess the ability of these circuits to dynamically switch payload expression in response to changes in TMP concentration, we assessed the reversibility of the circuits by incubating cells with or without TMP, switching to TMP-supplemented or no-TMP media after 24 hours, then measuring bioluminescence at 48 hours post- transfection (**Fig. 3e**). Both circuits could be effectively turned “on” by adding TMP and “off” by removing it, as indicated by changes in the bioluminescence signal, and the triple-DD circuit provided the most robust switching **(Fig. 3e)**.

### Regulated nsPs enable high-fidelity control of payload gene expression *in vivo*

Based on these promising *in vitro* results, we next evaluated the expression behavior of these engineered replicons *in vivo*. saRNAs encapsulated in LNPs and injected intramuscularly (i.m.) produced strong expression in mouse muscle fibers **(Supplementary Fig. S2)**. To regulate saRNA expression, we tested oral TMP delivery. Mass spectrometry showed that TMP levels in the blood could readily be tuned by adjusting its dose in animal chow (**Fig. 4a**). We selected 2 mg/g TMP as the high “on” state dose for our *in vivo* studies, as this dose achieves a plasma concentration of ∼0.5 µM, which is well above the 0.1 µM threshold where saturation is reached *in vitro*, ensuring robust regulation across biological variability.

**Figure 4:**
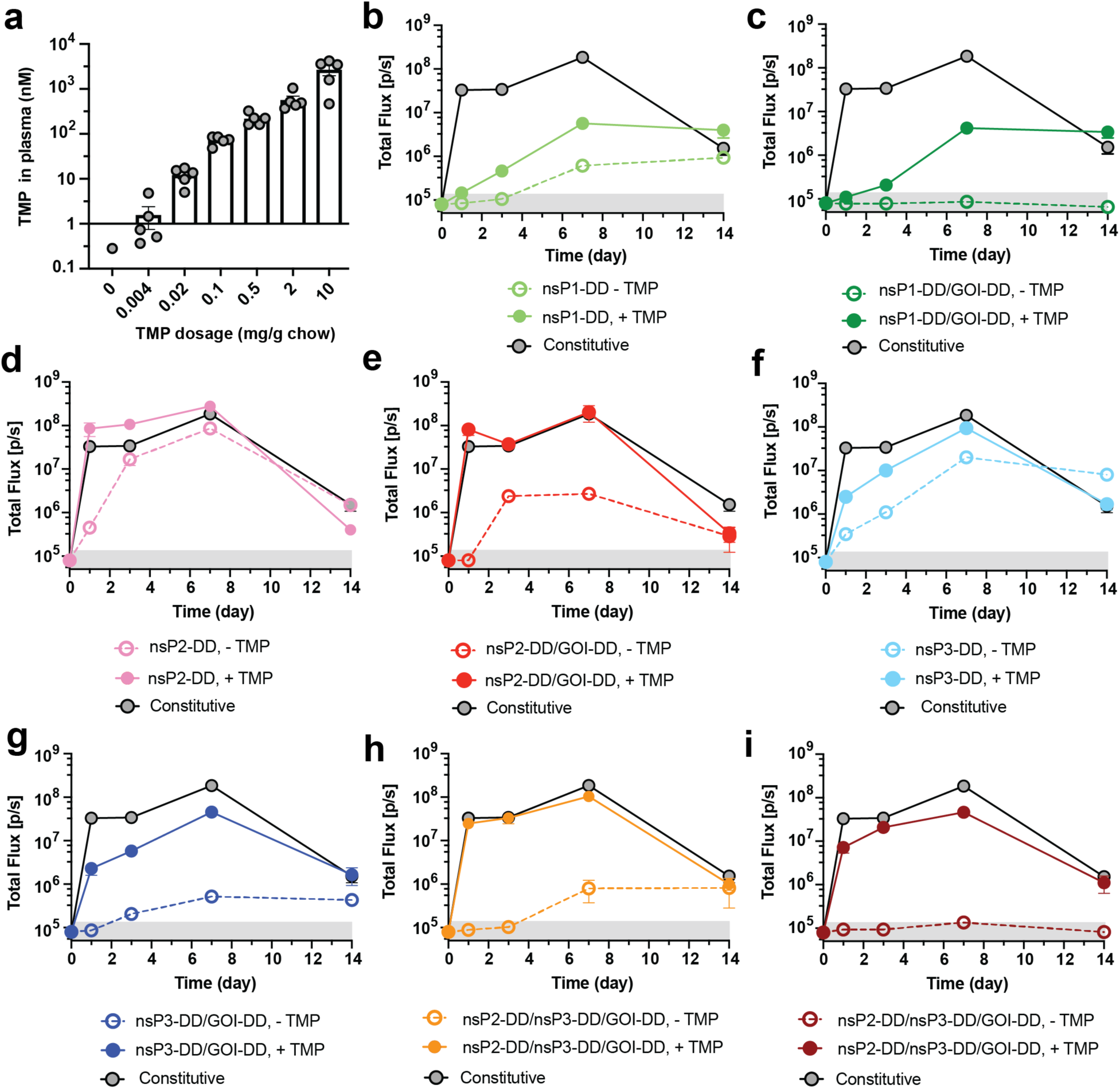
Testing of gene circuits differentiated muscle cells and *in vivo*. a, Plasma concentration of TMP achieved with different dosages of TMP. BALB/c mice were placed on diets with different dosages of TMP, and plasma samples were collected on day 3 post-TMP administration. Plasma TMP concentration was measured by LC-MS. **b-i,** Testing different circuits in mice. BALB/c mice were injected intramuscularly with fLuc-encoding saRNAs incorporating nsP1-DD **(b)**, nsP1-DD/GOI-DD **(c)**, nsP2-DD **(d)**, nsP2-DD/GOI-DD **(e)**, nsP3-DD **(f)**, nsP3-DD/GOI-DD **(g)**, nsP2-DD/nsP3-DD **(h)**, and nsP2-DD/nsP3-DD/GOI-DD **(i)** gene circuits. For groups receiving TMP, mice were provided with a TMP-supplemented diet with ad libitum access. FLuc expression was assessed at multiple time points post-injection using bioluminescence readings from an In Vivo Imaging System. The shaded gray area indicates background bioluminescence from naïve mice. Data are presented as mean ± SEM (*n* as indicated for panel **a** and 8-10 for panels **b-i**).

We administered regulated-saRNA circuits i.m. with or without oral TMP, and assessed payload expression longitudinally by bioluminescence imaging. Non-regulated saRNAs showed high fLuc expression within 1 day post-injection, maintained through day 7, with signal decaying by day 14 (**Fig. 4b**). Consistent with our *in vitro* findings, nsP1-DD saRNAs showed 10-fold lower peak expression in the "on" state, which was slow to reach steady-state and substantial “off’ state expression developing over time (**Fig. 4b**). Fusing DD to both nsP1 and the payload GOI reduced the off-state leakiness but did not improve the on-state expression level (**Fig. 4c**). The nsP2-DD circuit displayed rapid peak expression at levels comparable to constitutive saRNAs in the “on” state but also high background expression in the “off” state (**Fig. 4d**). Addition of DD to the payload lowered the “off” state background but it remained substantial (**Fig. 4e**). nsP3-DD and nsP3-DD/GOI-DD saRNAs showed expression behavior similar to the nsP2-DD constructs, but the nsP3-DD/GOI-DD saRNA had lower off-state expression and remained within the same order of magnitude as background levels throughout (**Fig. 4f-g**). Notably, as shown in **Fig. 4h-i**, the dual strategy of fusing DD to both nsP2 and nsP3 resulted in robust expression in the “on” state while ensuring minimal, non-leaky expression in the “off” state, especially when combined with DD fused to the payload GOI.

To determine the *in vivo* dose-response of TMP regulated-expression, we administered LNPs carrying nsP2- DD/nsP3-DD/GOI-DD saRNAs encoding fLuc i.m. in the presence of different doses of TMP and evaluated the fLuc bioluminescence signal after 2 days. FLuc expression was strongly TMP-dependent, with bioluminescence signals tunable over two orders of magnitude, achieving maximal expression at 0.5 mg/g oral TMP (**Fig. 5a**), and relative levels of expression were maintained over time (**Fig. 5b)**. The nsP3-DD/GOI-DD replicon also showed a high dynamic range of expression in the muscle in response to orally delivered TMP (**Extended Data** Fig. 5a). By contrast, a regulated circuit with DD only fused to the GOI showed significant leakiness and saturation at 0.1 mg/kg TMP on day 7, with an on:off bioluminescence signal ratio that did not exceed 10-fold **(Extended Data** Fig. 5b-c).

**Figure 5.**
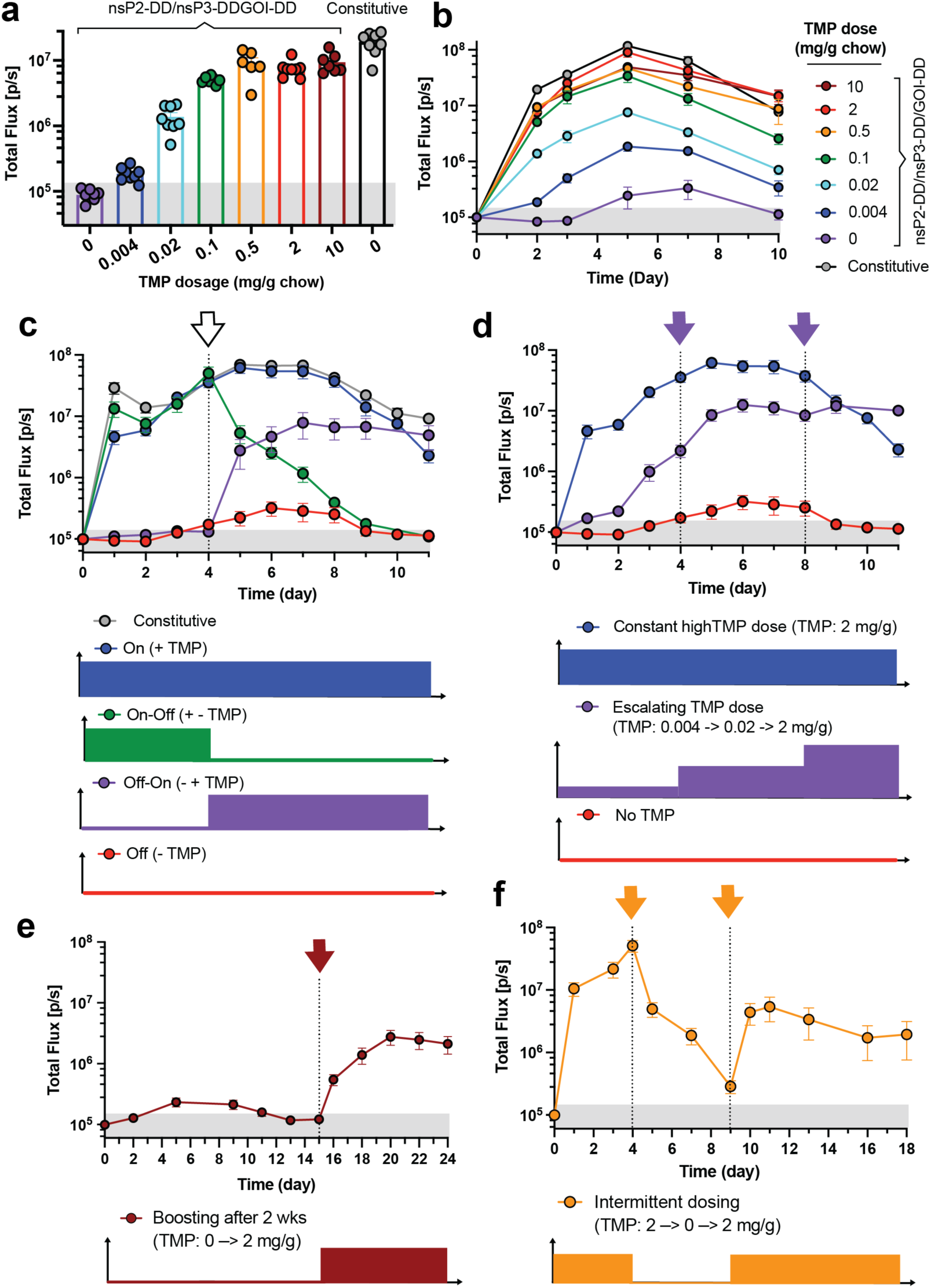
Controlling the kinetics of transgene expression from saRNA gene circuits by TMP. BALB/c mice were placed on diets supplemented with different dosages of TMP and injected intramuscularly with fLuc-encoding saRNAs incorporating the nsP2-DD/nsP3-DD/GOI-DD circuit. **a,b,** Adjusting the transgene expression level by changing the TMP dosage. Shown are **(a)** the bioluminescence signals at day 2 post saRNA injection and **(b)** longitudinally over 10 days. **c,** Reversibility of the nsP2-DD/nsP3-DD/GOI-DD saRNA circuit. The reversibility of the circuit was evaluated by switching mice from a no-TMP diet to a 2 mg/g chow diet on day 4 and vice versa. **d,e,f,** Achieving different transgene expression patterns with nsP2-DD/nsP3-DD/GOI-DD saRNA circuit. TMP dosages were adjusted at different time points to **(b)** escalate expression, **(e)** boost gene expression after 2 weeks, and **(f)** oscillate fLuc expression. The shaded gray area indicates background bioluminescence from naïve mice. Data are presented as mean ± SEM (*n* as indicated for panel **a** and 6-8 for panels **b-f**).

We next assessed dynamic switching of payload expression for the high-fidelity nsP2-DD/nsP3-DD/GOI-DD circuit. Transgene expression from non-regulated replicons showed an arc pattern, with high expression peaking around day 7 post-injection and then slowly decaying (**Fig. 5c**). By contrast, for the nsP-regulated saRNA, transitioning from a TMP-containing diet to a no-TMP diet on day 4 led to quenching of fLuc expression, approaching background levels over ∼5 days (**Fig. 5c**, green curve). Conversely, switching from a no-TMP to a TMP-containing diet led to a marked increase in gene expression within one day, plateauing within ∼3 days **(Fig. 5c**, purple curve). In addition, different TMP regimens, including escalating, “delayed on,” and oscillating doses, produced distinct expression profiles. Progressively increasing the TMP dose from 0.004 to 0.02 mg/g, to 2 mg/g, gave steadily increasing payload gene expression over the course of a week **(Fig. 5d**, purple curve). Administration of saRNA in the absence of TMP for an extended period of 15 days, followed by introduction of TMP allowed a delayed onset of expression peaking at day ∼20 **(Fig. 5e)**. Finally, alternating the TMP dose between 2 mg/g and 0 mg/g produced an oscillating pattern of gene expression **(Fig. 5f)**. The nsP3-DD/GOI-DD circuit showed similar dynamics but slower activation and lower initial "on" state levels compared to nsP2-DD/nsP3-DD/GOI-DD **(Extended Data** Fig. 5d-f).

We also tested the functionality of circuits in tumors in addition to muscles. nsP3-DD/fLuc-DD and nsP2-DD/nsP3- DD/fLuc-DD circuits were encapsulated in LNPs and injected intratumorally into subcutaneously implanted KP lung tumors. Mice were put on diets containing TMP dosages of 0, 0.02, or 2 mg/g chow. As expected, the bioluminescence signal was modulated by TMP dosage **(Extended Data** Fig. 6), demonstrating effective regulation of gene expression within the tumor environment. TMP-regulated saRNAs, therefore, enable reversible, dynamic control over transgene expression *in vivo*, providing a powerful tool for modulating gene expression in various biological systems.

### Using regulated saRNAs to augment B cell responses to vaccination

One application area of interest for high-fidelity small molecule-controlled gene expression is in the setting of vaccines, where the temporal pattern of antigen exposure can significantly impact the quality and magnitude of immune responses developed.^5,7,60^ To explore the potential of our optimal nsP2-DD/nsP3-DD/GOI-DD circuit in this setting, we employed it to express a clinically relevant HIV gp120-derived immunogen known as engineered outer domain-GT8 (eOD). We synthesized nsP2-DD/nsP3-DD saRNAs encoding eOD-DD as a secreted payload, and first evaluated eOD expression from replicon-transfected C2C12 cells. In the absence of TMP, no eOD was detectable, whereas in the presence of TMP, secreted eOD was detected at approximately 70 pg per 1000 transfected C2C12 cells, confirming that antigen expression was tightly controlled by TMP **(Fig. 6a)**.

**Figure 6.**
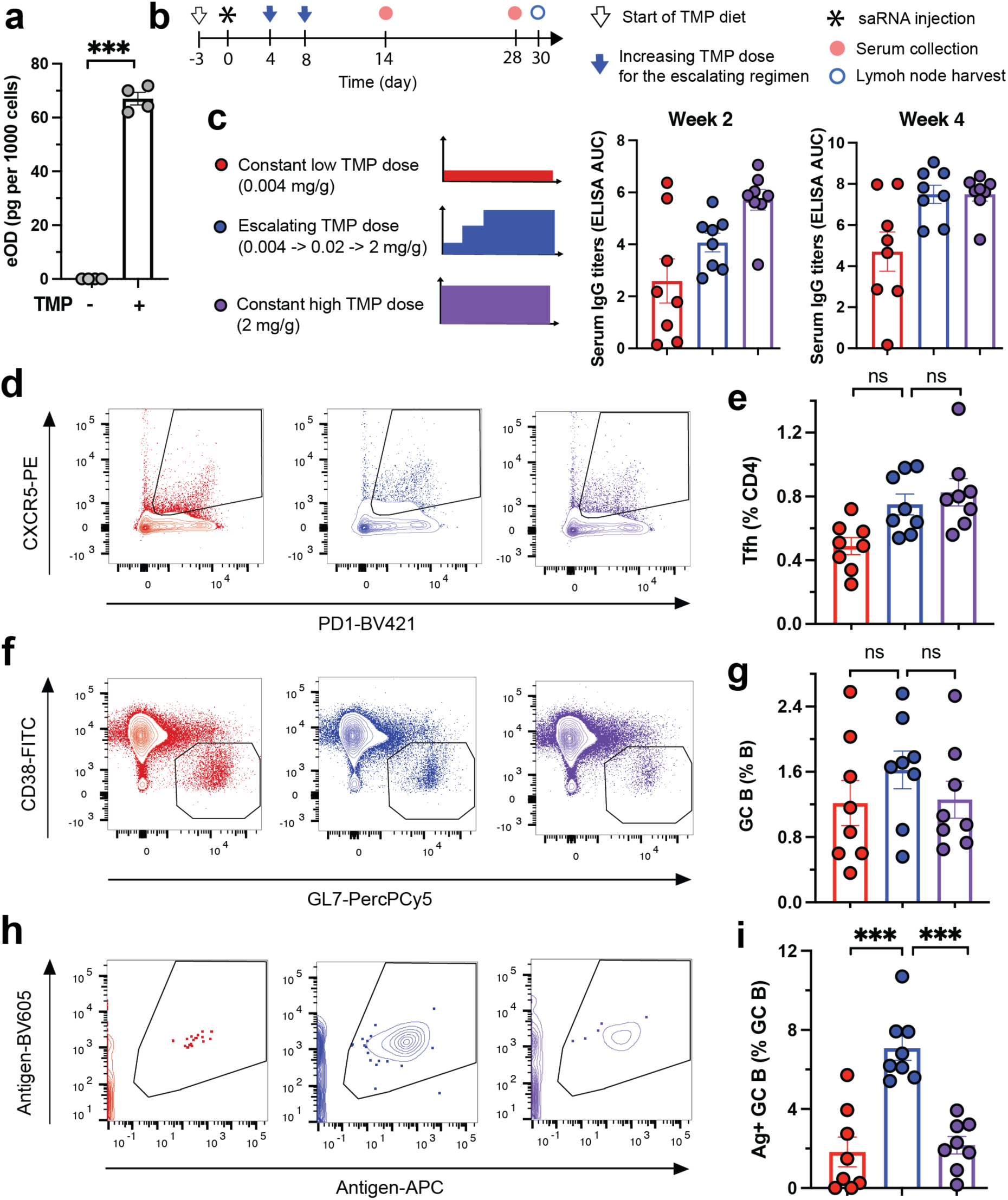
Controlling immune responses to nsP2-DD/nsP3-DD/GOI-DD saRNA vaccines using TMP. a, Regulation of eOD expression from saRNAs in vitro by TMP. C2C12 cells were transfected with a TMP-regulated eOD-encoding nsP2-DD/nsP3-DD/GOI-DD saRNA and incubated with or without TMP in the media. The eOD concentration in the cell supernatant was measured by ELISA. **b-i,** Immune responses to eOD-encoding nsP2- DD/nsP3-DD/GOI-DD saRNA vaccines under different TMP regimens. **(b)** Study diagram. BALB/c mice were immunized with eOD-encoding saRNAs encapsulated in lipid nanoparticles and placed on diets with specified TMP regimens. Antibody responses were quantified by ELISA at weeks 2 and 4 post-immunization. Germinal center responses were analyzed at week 4 post-immunization by collecting draining lymph nodes, staining for different markers, and performing flow cytometry. The panels show antibody responses **(c)**, representative flow cytometry plots and frequencies of Tfh cells (**d** and **e**, respectively), germinal center B cells (**f** and **g**, respectively), and antigen-specific GC B cells (**h** and **i**, respectively). Data are presented as mean ± SEM with indicated *n*. Statistical comparisons were performed using Student’s t test (A) or one-way ANOVA, followed by Tukey’s post-hoc test (e, g, and i). *, P < 0.05; **, P < 0.01; ***, P < 0.001.

We have previously shown that repeated protein vaccine injections with increasing concentrations every few days can greatly augment humoral immunity, especially germinal center (GC) responses that are critical for affinity maturation of rare B cells capable of generating protective antibodies.^61–63^ However such a dosing schedule is impractical for clinical implementation. We reasoned that small molecule-regulated saRNA could replicate this antigen exposure pattern through escalating oral TMP doses. Further, we hypothesized that GC responses to such an escalating pattern of antigen expression might outperform the near-constant high levels of antigen expression from traditional replicon immunization, by enforcing stricter selection on B cells through limited antigen availability early in the immune response.

To test this idea, mice were vaccinated i.m. with nsP2-DD/nsP3-DD/eOD-DD saRNA encapsulated in LNPs, and assigned to three TMP dose groups: low (0.004 mg/g chow), high (2 mg/g chow), and escalating (0.004 mg/g increasing to 0.02 mg/g after 4 days, then 2 mg/g after 4 more days, **Fig. 6b-c**). Antigen-specific IgG titers were significantly elevated in both the escalating and high TMP regimens compared to the low-TMP control group at both weeks 2 and 4 post-immunization. At week 2, titers were higher in the high TMP group than the escalating TMP regimen, but titers in these 2 groups equalized by week 4 **(Fig. 6c)**. However, more prominent effects were found when we examined GC responses primed by these different replicon expression regimens by flow cytometry **(Supplementary Fig. S3)**. The frequencies of GC B cells and follicular helper T cells (Tfh cells) were similar across all groups **(Fig. 6d-g)**. However, GC B cells that had acquired sufficient affinity to bind to fluorescent antigen probes were markedly enriched in the escalating group compared to the constant low or high TMP groups (**Fig. 6h-i**). These results suggest that escalating-expression immunization using regulated replicons enhances the maturation of antigen-specific B cells in the GC, a critical element for vaccines targeting difficult-to-neutralize pathogens where the GC must expand very rare precursor B cells capable of maturing into broadly neutralizing antibody-producing cells.

## Discussion

We show here that by combining the long-lived expression of saRNAs with small-molecule-based expression control, high fidelity dynamic gene expression tunable over multiple weeks can be achieved *in vitro* and *in vivo*. Integrating DDs with nsPs in the saRNA backbone presents a promising approach for RNA-level control of transgene expression by targeting saRNA amplification mechanisms. DD fusions to nsP1, nsP2, or nsP3 provided regulatory control of diverse payload proteins and enabled diverse transgene expression patterns. Importantly, the best-performing replicon designs functioned similarly across cells from different species and cell types, highlighting the robustness of this strategy.

We found significant differences in the effectiveness of the DD-fusion strategy based on which nsPs were engineered, reflecting the complex interplay of nsP functions in saRNA replication. Fusion to nsP1 significantly decreased on-state expression, likely due to interference with its function in RNA capping and replicase membrane association.^64–66^ nsP2 fusion allowed high on-state expression, but caused notable off-state leakiness, possibly due to nsP2’s multifunctional roles in transcriptional and translational shutoff in host cells and helicase and protease activities.^66^ The function of nsP3 is poorly understood, but it is thought to contribute to RNA synthesis and cytoplasmic granule formation.^66,67^ Fusion to nsP3 showed moderate expression in the "on" state with low leakiness in the "off" state. Interestingly, combining nsP2 and nsP3 fusions provided both high on-state expression and minimal off-state leakiness, suggesting a synergistic effect that enhances their collaborative function within the replication complex. Fusion to nsP4, the most conserved alphavirus protein, suppressed payload expression, indicating its RNA polymerase activity is likely impaired by DD fusion.^66^

Remarkably, TMP-dependent regulation persisted when regulated saRNAs were co-transfected with wildtype saRNAs. This likely results from distinct spherules forming during saRNA amplification,^68,69^ allowing autonomous function of regulated and constitutive saRNAs. This compartmentalization enables co-delivery of saRNAs carrying distinct payloads, where one is constitutively expressed, and the other is TMP-regulated. In the context of RNA vaccines, this could support simultaneous delivery of antigens and adjuvant cytokines, with the antigen being constitutively expressed while the adjuvant expression is temporally controlled, or vice versa, for enhanced immunity.

We demonstrated one potential application of regulated saRNA by using oral TMP delivery to program expression of an HIV antigen to escalate over the course of ∼1 week, enhancing antigen-specific B cell maturation in draining lymph nodes. We can also envision the potential of regulated replicons to enable prime-boost immunization strategies by pulsing antigen expression early and re-expressing a few weeks later via oral TMP delivery. Such drug-mediated boosting could reduce the need for repeat access to healthcare workers for immunization during epidemics and allow a needle-free vaccine boost. Future work could also explore incorporating non-antibiotic alternatives to TMP for stabilizing DHFR-based DDs to address concerns about antibiotic resistance;^70^ however, their lack of FDA approval may pose additional challenges for clinical translation.

In summary, we demonstrated an RNA-based gene circuit enabling precise control of transgene expression kinetics via an FDA-approved small molecule drug. This platform allows for high-fidelity control over gene expression, enabling gene expression to be regulated *in vivo* over several weeks. This expands the potential applications of saRNAs, as control of exposure to immunological payloads can further enhance immune responses, of interest for diverse therapeutic applications, including next-generation vaccines, gene therapies, and cancer treatments.

## Methods

### saRNA template DNA construction

A Venezuelan equine encephalitis (VEE) saRNA backbone, derived from the TC-83 strain, as described previously,^71^ was used for constructing saRNAs in this study. The saRNA vector was optimized for compatibility with Gibson Assembly, facilitating the efficient cloning of various constructs. Gene fragments encoding for the destabilizing domain (DD) and payload sequences were obtained through either as custom-ordered synthetic gene blocks from Integrated DNA Technologies (IDT) or via polymerase chain reaction (PCR) amplification of the saRNA open reading frame (ORF) sequences from sequence-verified plasmids. To enable the secretion of eOD and mCherry, a signal peptide from the spike glycoprotein of the Middle East respiratory syndrome-related coronavirus (MERS-CoV) was added to the N-terminus of the constructs. For transmembrane mCherry, the platelet-derived growth factor receptor beta (PDGFRβ) transmembrane domain was appended to the C-terminus, in addition to the signal peptide at the N-terminus. The Gibson Assembly method was employed to clone the DD and payload gene fragments into the VEE saRNA backbone. The plasmid DNA was extracted and purified from bacterial cultures using the QIAGEN Midi Kit. Bacteria harboring the plasmid of interest were grown in LB medium containing the appropriate antibiotic for plasmid selection. Following bacterial growth, plasmid extraction was performed according to the manufacturer’s protocol, which includes cell lysis, DNA binding, washing, and elution steps to obtain high-purity plasmid DNA.

### saRNA synthesis

To synthesize saRNA RNA, template plasmid DNA was linearized via endonuclease digestion and purified using PureLink PCR Purification columns (ThermoFisher #K310002) according to the manufacturer’s instructions. Subsequently, 20 μL *in vitro* transcription (IVT) reactions were performed using the HiScribe T7 High Yield RNA Synthesis Kit (NEB #E2040S) with 1-2μg of linear DNA template, scaled as needed. The IVT product was purified using PureLink RNA Mini columns (ThermoFisher #12183018A) according to the manufacturer’s instructions. The RNA was then capped and methylated using the ScriptCap Cap 1 Capping System (CellScript #C-SCCS2250), following the manufacturer’s instructions. After capping and methylation, the RNA was purified a final time using PureLink RNA Mini columns. The quality of the resulting saRNAs was assessed using UV-Vis spectrophotometry and gel electrophoresis.

### Animal maintenance

Female BALB/c mice (JAX Stock No. 000651), aged 6–8 weeks, were housed in the animal facility at the Massachusetts Institute of Technology (MIT) under a 12-hour light-dark cycle. All animal experiments and procedures were conducted in accordance with federal, state, and local regulations, under an IACUC-approved protocol (MIT Committee on Animal Care (CAC) Protocol Number: 2303-000-488). Trimethoprim (TMP)- supplemented diets were purchased from Inotiv. These diets were provided based on experimental requirements, with the mice having ad libitum access to TMP-containing chow for the indicated durations. The concentration of TMP in the diet and the timing of administration were chosen according to the experimental design.

### Cell culture and transfection

RNA transfections were performed using C2C12 mouse myoblasts, KP lung cancer cells, BHK-21 cells, and human HEK293T cells. C2C12, BHK-21 and HEK293T cell lines were purchased from ATCC. KP cell line was generously provided by Dr. Tyler Jacks (Massachusetts Institute of Technology, Cambridge). BHK-21 cells were cultured in Eagle’s Minimum Essential Medium (EMEM, ATCC), C2C12, KP, and HEK293T cells were grown in Dulbecco’s Modified Eagle Medium (DMEM, ATCC). All media were supplemented with 10% 10% FBS (Sigma-Aldrich) and 1% Penicillin-Streptomycin (100 U/mL penicillin, 100 µg/mL streptomycin; Thermo Fisher Scientific),

and maintained at 37 °C in a 5% CO2 atmosphere. Cells nearing 70% confluence were electroporated using the Neon Transfection System (Life Technologies), with settings optimized for each cell line: 1,100 mV, 40 ms, and 1 pulse for BHK-21 cells; 1,400 mV, 20 ms, and 1 pulse for C2C12 cells; 1,005 mV, 35 ms, and 2 pulses for KP cells; 1,200 mV, 10 ms, and 3 pulses for HEK293T cells.

For transfections, ∼5x10^6^ C2C12 cells were used per 100 μL in a single well of a 24-well plate (Corning). The amount of RNA transfected was 1 μg per 100,000 BHK-21 cells and 100 ng per 50,000 C2C12 cells, unless specified otherwise. The experiments were conducted in either 24- or 96-well plates (Corning) with adjusted plating densities. Stock solutions of 2 mM TMP in PBS were diluted to final concentrations of 10 μM.

To differentiate C2C12 cells into myotubes, cells were transferred to differentiation medium containing 2% horse serum without FBS and incubated for 5-7 days. Cells were subsequently transfected with saRNAs encapsulated in LNPs.

### Evaluation of *in vitro* transfection

For experiments using firefly luciferase (fLuc) as the reporter cargo, transfected cells were cultured in black 96- well plates with TC-treated bottoms. To measure luminescence, the supernatant was removed and replaced with 100 μL of fresh media, followed by the addition of 100 μL luciferin at 15 mg/mL. Bioluminescence was measured 10 minutes after the addition of luciferin.

For experiments using mCherry as the reporter, transfected cells were plated in 24-well plates and harvested as follows: cells were washed with PBS (Corning), trypsinized (Corning), quenched with cell growth media, and resuspended in PBS. After staining with Zombie Aqua, cells were washed with FACS buffer (PBS + 2% FBS + 2.5 mM EDTA) and resuspended in FACS buffer. Flow cytometry was performed using a BD FACSCelesta or FACSSymphony A1 Flow Cytometer System (BD Biosciences) equipped with 405, 488, and 561 nm lasers. Initial data collection was done using FACSDiva software, and flow cytometry data analysis was performed using FlowJo software. For differentiating C2C12 cells into myofibers, C2C12 cells were cultured in DMEM media supplemented with 2% horse serum for 6 days. Transfection of myofibers was performed by encapsulating RNAs in lipid nanoparticles as described below. Cell viability was assessed at indicated time points post-transfection using the CellTiter-Glo™ Luminescent Cell Viability Assay (Promega) according to the manufacturer’s instructions. Payload expression was measured by ELISA using an mCherry ELISA Kit (Abcam) according to the manufacturer’s protocol, and bioluminescence measurement for fLuc.

### Lipid nanoparticle synthesis

For *in vivo* transfection and *in vitro* transfection of differentiated C2C12 myofibers, saRNAs were encapsulated in lipid nanoparticles using the Precision Nanosystems Ignite microfluidic mixing platform. The RNA phase was prepared in 10 mM citrate buffer at pH 3.0 (CAT#J61391-AK; Alfa Aesar) at a concentration of 60 µg/ml. Lipids were prepared in ethanol and consisted of N1,N3,N5-tris(3-(didodecylamino)propyl)benzene-1,3,5-tricarboxamide (TT3, synthesized as described previously^72^), cholesterol (Chol, Avanti Polar Lipids), 1,2-dioleoyl-sn-glycero-3- phosphoethanolamine (DOPE, Avanti Polar Lipids), and 1,2-dimyristoyl-sn-glycero-3-phosphoethanolamine-N- [methoxy(polyethylene glycol)-2000 (DMPE-PEG, Avanti Polar Lipids) at a 22:33:44:1 TT3/DOPE/Chol/DMPE-PEG molar ratio. Both phases were loaded into a syringe (BD) and injected into the NxGen microfluidic cartridge of a NanoAssemblr Ignite instrument (Precision Nanosystems) set to a total flow rate of 12 mL/min and 2:1 RNA:lipid flow rate ratio. The resulting LNPs were then dialyzed against RNase-free PBS using 20 K MWCO Slide- A-Lyzer™ MINI Dialysis cassettes (ThermoFisher Scientific) at 25°C with a 45-minute initial dialysis and a subsequent 75-minute dialysis after a buffer exchange. The final saRNA-LNPs were diluted in PBS to achieve the required concentrations for *in vivo* injection.

### *In vivo* evaluation of gene circuits in mice

To assess the transgene expression from different gene circuits in response to TMP, saRNAs encoding fLuc as a reporter were injected into BALB/c mice. Each mouse received 1 μg of RNA-loaded LNPs via intramuscular injection into the left and right gastrocnemius muscles. At various time points post-injection, the mice were subcutaneously administered 200 μL of luciferin (15 mg/mL in PBS) and imaged using the In Vivo Imaging System (Xenogen IVIS 200; PerkinElmer) 10 minutes after luciferin injection. Bioluminescence imaging data were analyzed using Living Image software (PerkinElmer). Regions of interest (ROIs) were drawn around the injection sites in both left and right gastrocnemius muscles, and the total flux (photons/second) within each ROI was measured to quantify fLuc expression.

### *In vivo* immunization

BALB/c mice were immunized via intramuscular injection with 0.4 μg of LNP-encapsulated saRNA RNA in each gastrocnemius muscle. Prior to vaccination, mice were assigned to different treatment groups and fed diets supplemented with varying concentrations of TMP: no TMP, 0.004 mg/g TMP, or 2 mg/g TMP. The diet regimen began three days before immunization. For the escalating treatment group, TMP concentration was incrementally increased from 0.004 mg/g to 0.01 mg/g on day 5, and further to 2 mg/g on day 10 post-immunization. Four weeks post-vaccination, blood was collected via retro-orbital bleeding into Z-gel PP tubes (Sarstedt, CAT#41.1500.005). Serum was isolated by centrifugation at 10,000 ×g for 5 minutes.

### Assessing antibody responses

Antibody responses to immunization were measured by Enzyme-Linked Immunosorbent Assay (ELISA). NUNC MaxiSorp plates were coated with 1 μg/mL purified eOD immunogen in PBS overnight at 4°C, followed by blocking with 10% BSA in PBS overnight at 4°C. Serum samples were initially diluted 20-fold in blocking buffer, then serially diluted 4-fold. Diluted sera were incubated on the blocked plates for 2 hours. HRP-conjugated immunoglobulin (Bio-Rad) was used as detection antibody at a 1:5000 dilution. The mouse VRC01 monoclonal antibody, which targets the HIV envelope CD4 binding site was used as a positive control. Signal development was achieved using 3,3′,5,5′-Tetramethylbenzidine (TMB) substrate, and absorbance was measured at 450 nm and 540 nm using a microplate reader. The final signal was calculated by subtracting the 540 nm reading from the 450 nm reading. To quantify antibody responses, the Area Under the Curve (AUC) of the final signal versus log-transformed sample dilution was calculated using GraphPad Prism software (Version 10.2.2).

### Germinal center assay

To assess the germinal center (GC) response following immunization, draining lymph nodes (popliteal and iliac) were collected and mechanically dissociated by passing them through 70 µm strainers to obtain single-cell suspensions. Zombie Aqua (BioLegend) in PBS was used to stain for cell viability. Antibody staining was performed in flow cytometry buffer (PBS, 2% FBS, and 2.5 mM EDTA). To prepare tetramer probes, biotinylated eOD antigen was conjugated with either streptavidin-BV605 or streptavidin-APC (both from BioLegend) and incubated for 30 minutes. Cells were stained with Zombie Aqua Fixable Viability Dye (BioLegend) for 15 minutes at room temperature, washed in flow cytometry buffer, and then incubated with 20 nM diluted tetramer probes for 30 minutes to detect antigen-specific B cells. Fc-mediated binding was blocked with anti-CD16/32 (2.5 μg/mL; 93; BioLegend) at 4°C for 10 minutes, followed by the addition of antibodies for cell surface staining at 4°C for 30 minutes. The following antibodies were used for staining: anti-GL7 PerCPCy5.5 (clone GL7; BioLegend), anti- CD38 AF488 (clone 90; BioLegend), anti-B220 PECy7 (clone RA3-6B2; BioLegend), and anti-CD4 BV711 (clone GK1.5; BioLegend), anti-B220 PECy7 (clone RA3-6B2; BioLegend), anti-CXCR5 PE (clone L138D7; BioLegend), and anti-PD1 BV421 (clone 29F.1A12; BioLegend). Finally, the stained cells were fixed with 4% paraformaldehyde (ThermoFisher Scientific) for 15 minutes at room temperature, washed, and resuspended in FACS buffer for flow cytometric analysis on a BD LSR Fortessa flow cytometer (BD Biosciences). Cells were gated based on forward and side scatter properties to exclude debris and doublets. Live cells were identified using the Zombie Aqua viability dye. GC B cells were defined as B220+CD4-GL7+CD38lo, and follicular helper T cells were identified as B220-CD4+CXCR5+PD1+.

### RNA extraction and qPCR

RNA was extracted from cells transfected with saRNAs to quantify the relative amounts of saRNA backbone and subgenomic RNAs. Cells were trypsinized, and the trypsinization was quenched with media, then collected and centrifuged. The resulting cell pellet was resuspended in RLT lysis buffer (Qiagen) containing 0.1% β-mercaptoethanol. RNA extraction was performed using the Chemegic 360 system with the chemagic RNA Tissue 10mg Kit H96 (Revvity) according to the manufacturer’s protocol. RNA integrity was assessed using an Agilent 2100 Bioanalyzer, and samples with an RNA Quality Number (RQN) less than 6 were excluded. cDNA was synthesized using the iScript™ Reverse Transcription Supermix (BioRad) following the manufacturer’s protocol and using a SimpliAmp Thermal Cycler (Applied Biosystems). The resulting cDNA was subjected to qPCR using iTaq Univer SYBR Green Supermix (BioRAD) on a QuantStudio 3 Real-Time PCR Instrument. For primer design targeting the saRNA positive strand, negative strand, and fLuc, five sets of primers were initially selected and screened. Primers producing more than one melting peak or non-linear Cq vs. cDNA dilution were excluded.

Mouse GAPDH primers were from IDT (Forward: catalog# 51-01-07-12; Reverse: 51-01-07-13). The relative amounts of each target were normalized to GAPDH on a per-sample basis.

Primer sequences:

Positive strand nsP3 Forward: TTCCCGGAAAGCTTCGATTTA Positive strand nsP3 Reverse: TCACCTTCAACCTCCGAAAC fLuc Forward: CACGCACGAGATCCCATATT fLuc Reverse: GCGGAACCCACATATCAAGTA

### Model fitting and predictions

4PL model fitting was performed using the dr4pl R package.^73^ The 4PL model is represented by the equation:

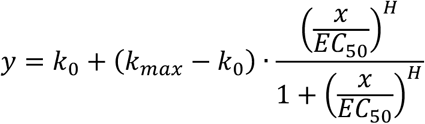

where x and y denote the TMP dose and resulting fLuc expression level, respectively. 𝑘_!_and 𝑘_"#$_represent the minimum and maximum asymptotes, respectively, 𝐸𝐶_%!_ is the inflection point, and 𝐻 is the slope at the inflection point. Assuming multiplicative effects of circuit compositions, we predicted fLuc expression levels in nsP-DD/fLuc- DD circuits by multiplying the fitted fold changes of individual parts with the maximum fLuc expression level from the circuit. The goodness-of-fit of our models, both for fitted and predicted curves, is evaluated using R-squared.

R-squared is one minus the quotient of summed squares of residuals by variance, i.e., [inline, where 𝑦, 𝑦(, 𝑦) represent log2-transformed observed fLuc levels, log2-transformed fitted/predicted fLuc levels, and average log2- transformed observed fLuc levels, respectively.

### Statistics

All graphs were generated using GraphPad Prism Version 10.2.2, and statistical analyses were conducted within the same software. Statistical comparisons were conducted using ANOVA, followed by the appropriate post hoc test as detailed in the figure legends. Statistical significance was defined at thresholds of *p < 0.05, **p < 0.01, and ***p < 0.001.

## Reporting Summary

Further information on research design is available in the Nature Research Reporting Summary linked to this article.

## Data Availability

The main data supporting the results in this study are included within the paper and its Supplementary Information. Any additional data supporting the findings of this study are also available from the corresponding authors on reasonable request. Source data will be provided with this paper.

## Acknowledgements

This work was supported in part by the Koch Institute Support (core) Grant 5P30-CA014051, award CA265706 (to DJI and RW), and awards UM1AI144462, AI175489 and AI125068 (to DJI) from the NIH. We thank the Koch Institute’s Robert A. Swanson (1969) Biotechnology Center for technical support, specifically the Flow Cytometry, Genomics, Media Preparation Core, Biopolymers & Proteomics Core, and the Preclinical Modeling, Imaging & Testing (PMIT) Core Facilities. Parisa Yousefpour is supported by an NRSA F32 fellowship (award AI164829) from the NIH. DJI is an investigator of the Howard Hughes Medical Institute. We also thank Dr. William R. Schief from the Scripps Research Institute for providing the sequence for the HIV immunogen eOD used in this study.

## Author Contributions

PY, RW, and DJI conceived the study, outlined experiments, interpreted the results, and wrote the manuscript. PY conducted in vitro circuit screening, testing, and in vivo mouse studies. JL and YL contributed to saRNA circuit design and testing. JW performed the model fitting and prediction. TR, JD, and LR performed in vitro transcription for saRNA synthesis. JG and KS contributed to the design and execution of experiments and data analysis. AL1, KQ, AL2, RY, and MG assisted with executing experiments. YZ synthesized the TT3 ionizable lipid. All authors reviewed and revised the manuscript. YD, RW, and DJI supervised the project.

## Extended Data

**Extended Data Figure 1:**
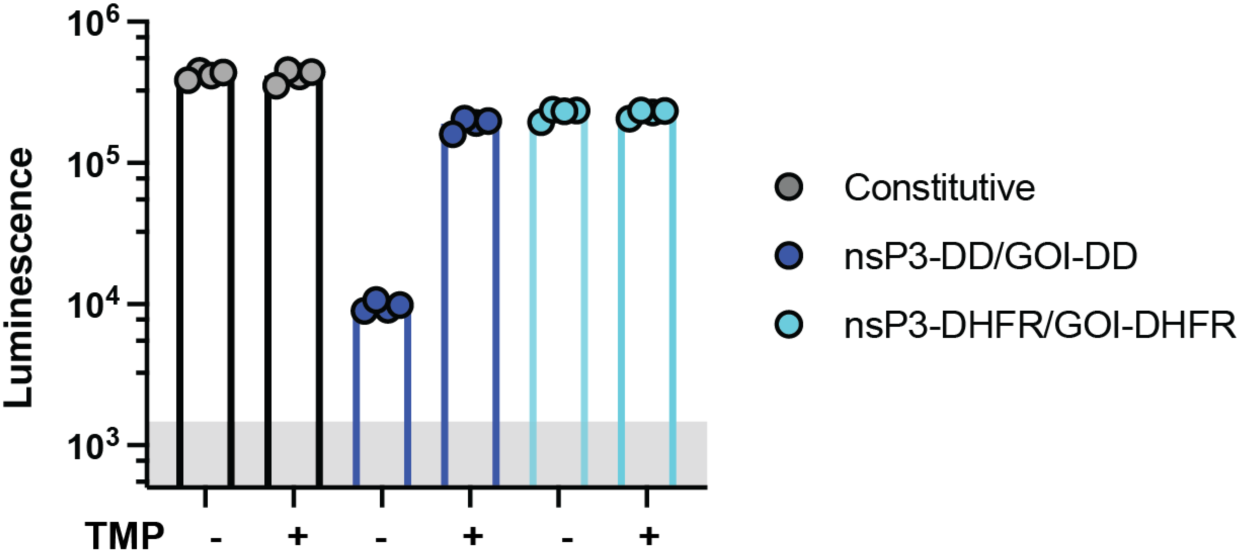
Impact of wild-type DHFR versus the mutated DHFR (DD) on transgene expression in response to TMP. C2C12 cells were electroporated with saRNAs where DD or DHFR was fused to nsp3 and fLuc, then incubated with or without TMP. Transgene (fLuc) expression was evaluated after 24 hours by measuring bioluminescence. Data are presented as mean ± SEM with indicated *n*.

**Extended Data Figure 2:**
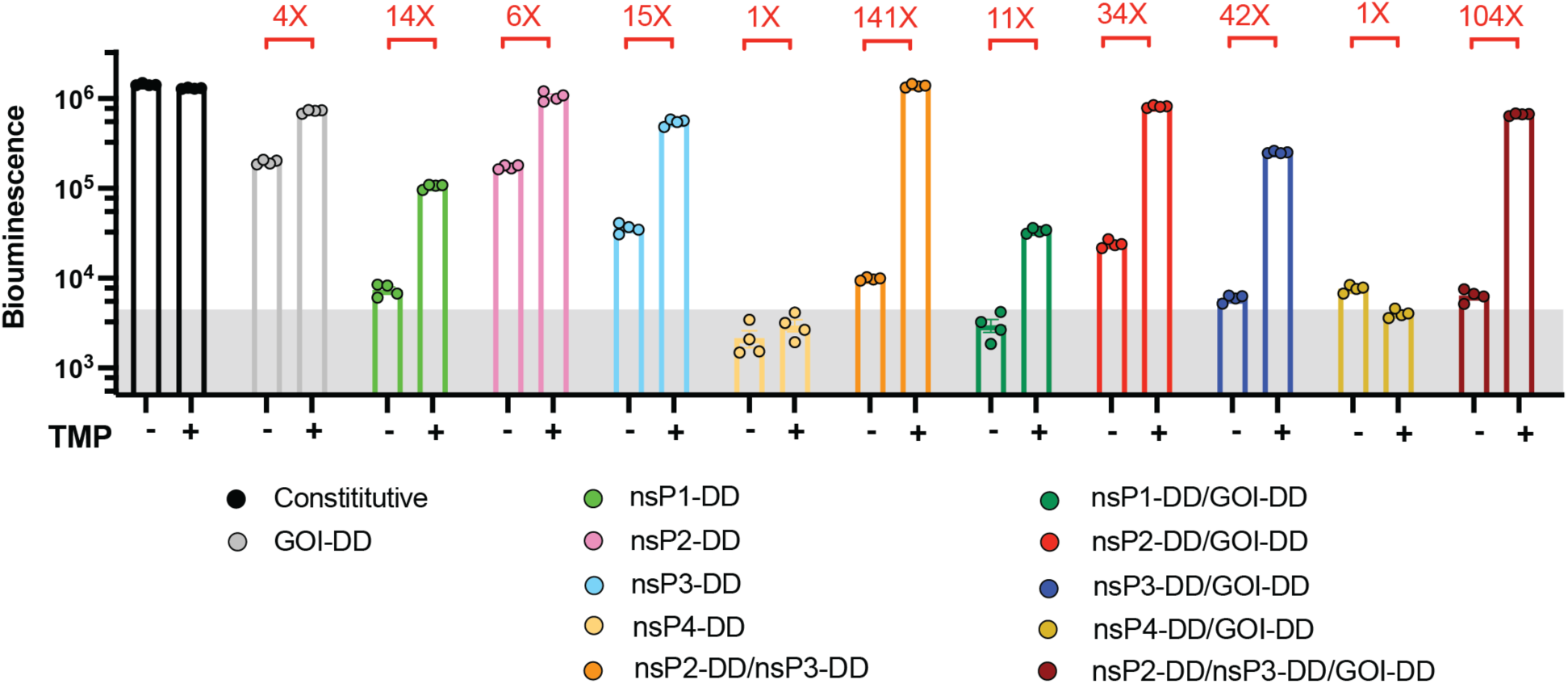
Regulation of transgene expression from different DD-based saRNA circuits in response to TMP in human HEK293T cells. Human embryonic kidney (HEK293T) cells were transfected with different saRNA gene circuits encoding fLuc and were cultured with or without TMP post-transfection. FLuc expression was assessed by bioluminescence at 24 hours post-transfection. Data are presented as mean ± SEM with indicated *n*.

**Extended Data Figure 3:**
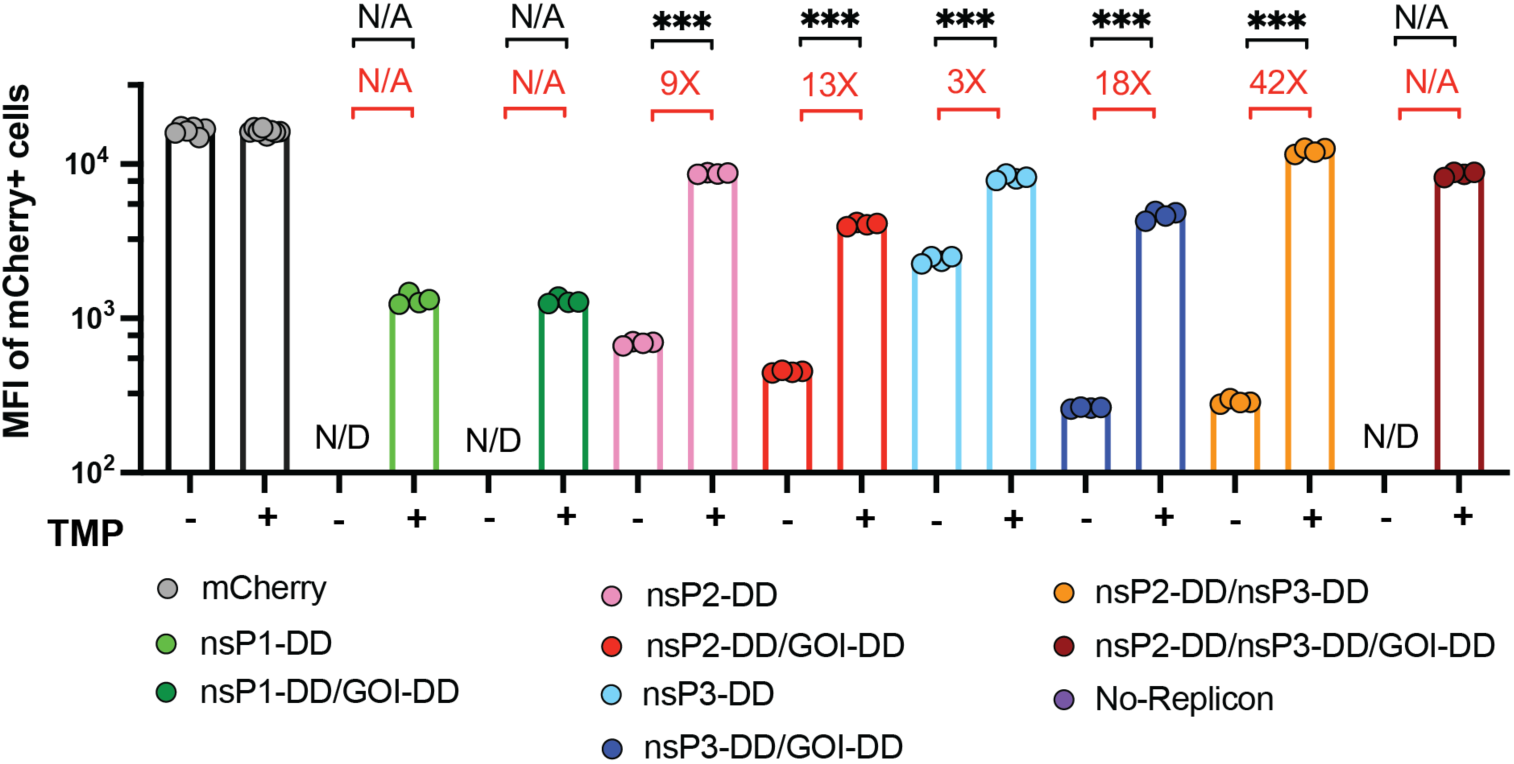
Levels of transgene expression from regulated saRNA circuits. C2C12 cells were transfected with saRNA gene circuits encoding mCherry. At 24 hours post-transfection, gene expression was evaluated at the individual cell level by flow cytometry. Shown is the median fluorescence intensity (MFI) of mCherry+ cells. N/D: Not Detected, indicating no or too few mCherry+ cells to reliably calculate MFI. Data are presented as mean ± SEM with indicated *n*. Statistical comparisons were performed using one-way ANOVA, followed by Tukey’s post-hoc test. ***, P < 0.001.

**Extended Data Figure 4:**
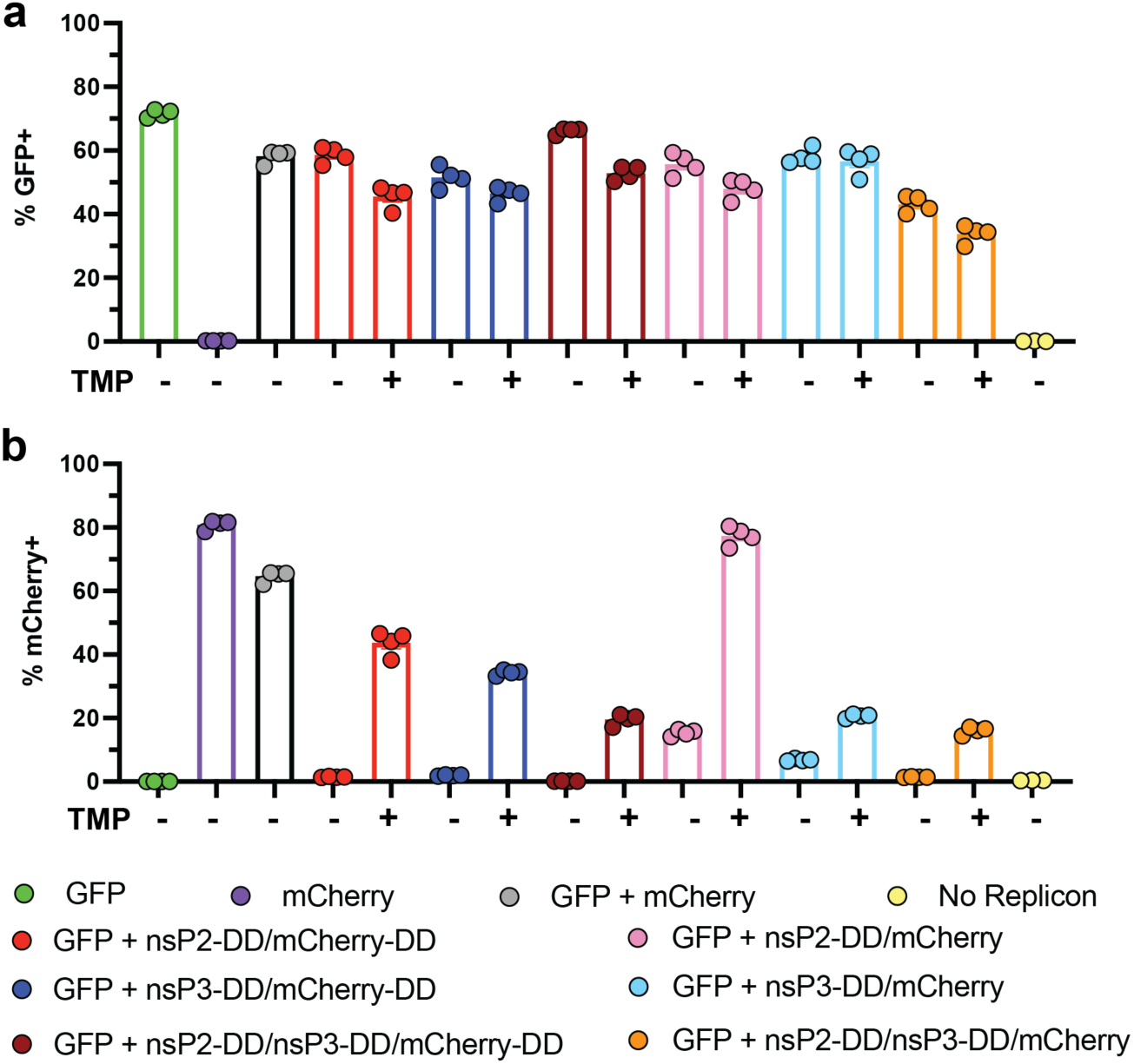
Co-transfection with constitutive and TMP-regulated saRNAs. a, b, C2C12 cells were electroporated with a 1:1 mixture of constitutive GFP-encoding saRNA and TMP-regulated mCherry- encoding saRNAs. Cells were incubated with or without TMP, harvested 24 hours post-transfection, harvested 24 hours post-transfection, and assessed for GFP **(a)** and mCherry **(b)** expression by flow cytometry. Data are presented as mean ± SEM with indicated *n*.

**Extended Data Figure 5:**
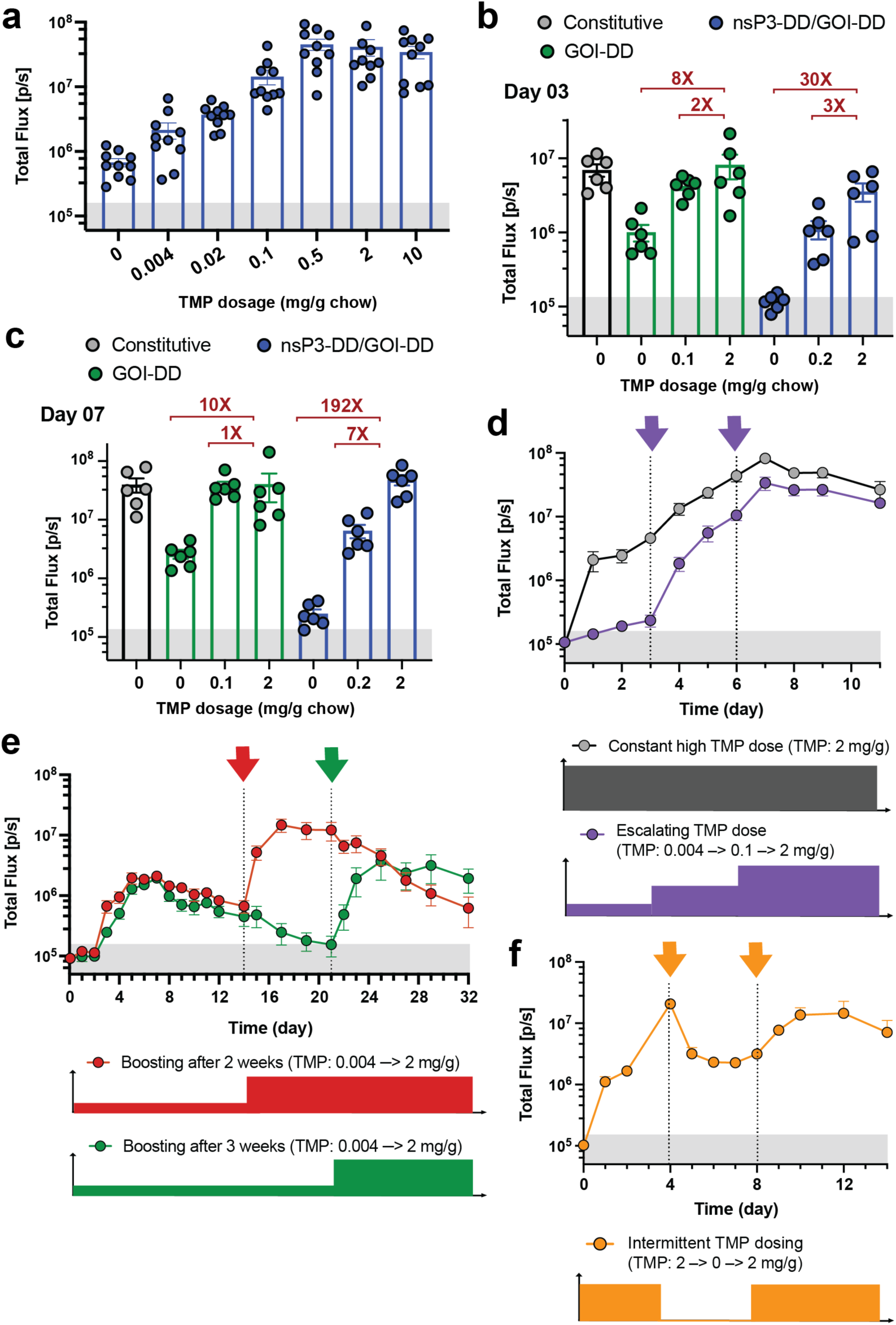
Controlling the kinetics of gene expression from nsP3-DD/GOI-DD and GOI-DD saRNA circuits. a, Adjusting the transgene expression level by changing the TMP dosage. BALB/c mice were placed on diets supplemented with different dosages of TMP and injected intramuscularly with fLuc-encoding saRNAs incorporating the nsP3-DD/GOI-DD circuit. FLuc expression was measured by bioluminescence imaging. **b,c,** Comparison of replication-controlled and non-replication-controlled TMP-responsive saRNAs. Mice were placed on diets with varying TMP dosages, and received intramuscular injections of saRNAs containing either the nsP3-DD/GOI-DD circuit (replication-controlled) or the GOI-DD circuit (non-replication-controlled). Following the injections, fLuc expression was monitored via bioluminescence imaging at day 3 **(b)** and 7 **(c)** post-injection. **d,e,f,** Achieving different transgene expression patterns. TMP dosages were adjusted at different time points to **(d)** escalate expression, **(e)** boost expression after 2 or 3 weeks, and **(f)** oscillate expression. FLuc expression was measured by bioluminescence imaging. Data are presented as mean ± SEM (*n* as indicated for panels **a-c** and 6-8 for panels **d-f**).

**Extended Data Figure 6:**
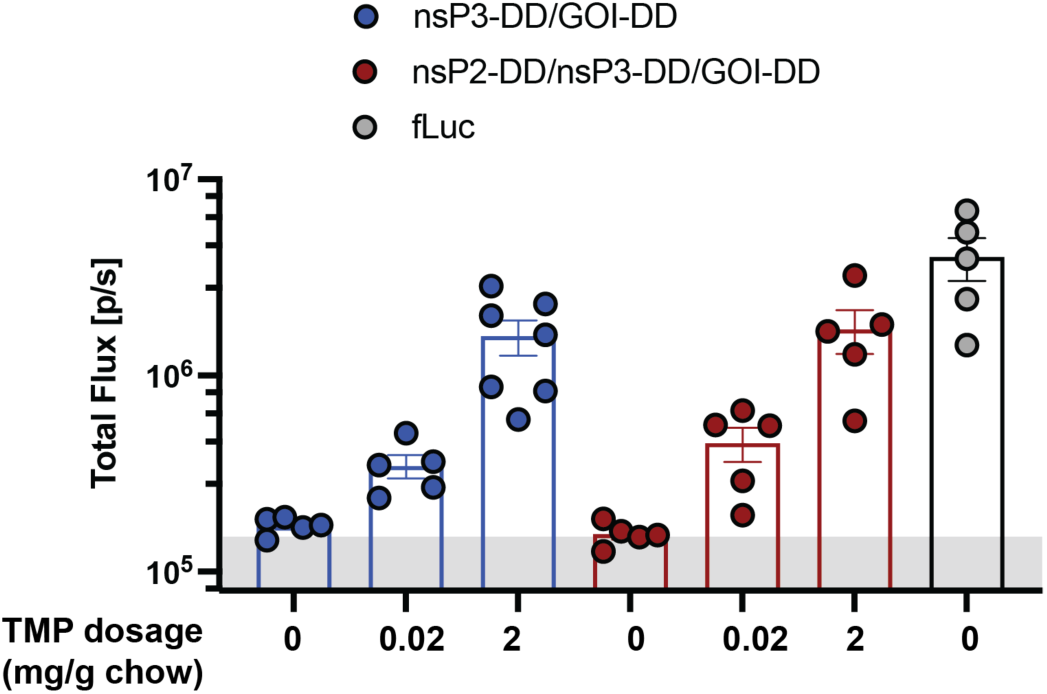
In vivo functionality of saRNA circuits in tumors. saRNAs encoding fLuc and incorporating nsP3-DD/GOI-DD or nsP2-DD/nsP3-DD/GOI-DD circuits were encapsulated in LNPs and injected intratumorally into subcutaneously implanted KP tumors. Mice were fed diets containing different dosages of TMP, and bioluminescence signals were quantified at day 2 post-injection using IVIS. Data are presented as mean ± SEM with indicated *n*.

## References

1. Rothschilds, A. et al. Order of administration of combination cytokine therapies can decouple toxicity from efficacy in syngeneic mouse tumor models. OncoImmunology 8, e1558678 (2019).

2. Messenheimer, D. J. et al. Timing of PD-1 blockade is critical to effective combination immunotherapy with anti-OX40. Clin. Cancer Res. 23, 6165–6177 (2017).

3. Chiocca, E. A. et al. Regulatable interleukin-12 gene therapy in patients with recurrent high-grade glioma: Results of a phase 1 trial. Sci. Transl. Med. 11, (2019).

4. Zhang, P. et al. Phase I trial of inducible caspase 9 T cells in adult stem cell transplant demonstrates massive clonotypic proliferative potential and long-term persistence of transgenic T cells. Clin. Cancer Res. 25, 1749– 1755 (2019).

5. Johansen, P. et al. Antigen kinetics determines immune reactivity. Proc. Natl. Acad. Sci. 105, 5189–5194 (2008).

6. Baden, L. R. et al. Timing of plasmid cytokine (IL-2/Ig) administration affects HIV-1 vaccine immunogenicity in HIV-seronegative subjects. J. Infect. Dis. 204, 1541–1549 (2011).

7. Tam, H. H., et al. Sustained antigen availability during germinal center initiation enhances antibody responses to vaccination. Proc. Natl. Acad. Sci. 113, E6639–E6648 (2016).

8. Bulcha, J. T., Wang, Y., Ma, H., Tai, P. W. L. & Gao, G. Viral vector platforms within the gene therapy landscape. Signal Transduct. Target. Ther. 6, 53 (2021).

9. Shirley, J. L., Jong, Y. P. de, Terhorst, C. & Herzog, R. W. Immune responses to viral gene therapy vectors. Mol. Ther. 28, 709–722 (2020).

10. Wang, J.-H., Gessler, D. J., Zhan, W., Gallagher, T. L. & Gao, G. Adeno-associated virus as a delivery vector for gene therapy of human diseases. Signal Transduct. Target. Ther. 9, 78 (2024).

11. Ronzitti, G., Gross, D.-A. & Mingozzi, F. Human immune responses to adeno-associated virus (AAV) vectors. Front. Immunol. 11, 670 (2020).

12. Poletti, V. & Mavilio, F. Designing lentiviral vectors for gene therapy of genetic diseases. Viruses 13, 1526 (2021).

13. Sabatino, D. E. et al. Evaluating the state of the science for adeno-associated virus integration: An integrated perspective. Mol. Ther. 30, 2646–2663 (2022).

14. Russell, D. W. AAV Vectors, Insertional Mutagenesis, and Cancer. Mol. Ther. 15, 1740–1743 (2007).

15. Costello, A. et al. Leaky expression of the TET-On system hinders control of endogenous miRNA abundance. Biotechnol J 14, e1800219 (2019).

16. Meyer-Ficca, M. L. et al. Comparative analysis of inducible expression systems in transient transfection studies. Anal. Biochem. 334, 9–19 (2004).

17. Mizuguchi, H. & Hayakawa, T. Characteristics of adenovirus-mediated tetracycline-controllable expression system. 1568, 21–29 (2001).

18. Zhuo, C. et al. Spatiotemporal control of CRISPR/Cas9 gene editing. Signal Transduct. Target. Ther. 6, 238 (2021).

19. Kampmann, M. CRISPRi and CRISPRa screens in mammalian cells for precision biology and medicine. ACS Chemical Biology 13, 406–416 (2018).

20. Kolos, J. M., Voll, A. M., Bauder, M. & Hausch, F. FKBP ligands—where we are and where to go? Front. Pharmacol. 9, 1425 (2018).

21. Goverdhana, S. et al. Regulatable gene expression systems for gene therapy applications: progress and future challenges. Mol. Ther. 12, 189–211 (2005).

22. Hou, X., Zaks, T., Langer, R. & Dong, Y. Lipid nanoparticles for mRNA delivery. Nat. Rev. Mater. 6, 1078– 1094 (2021).

23. Aliahmad, P., Miyake-Stoner, S. J., Geall, A. J. & Wang, N. S. Next generation self-replicating RNA vectors for vaccines and immunotherapies. Cancer Gene Ther. 30, 785–793 (2023).

24. Comes, J. D. G., Pijlman, G. P. & Hick, T. A. H. Rise of the RNA machines – self-amplification in mRNA vaccine design. Trends Biotechnol. 41, 1417–1429 (2023).

25. Dailey, G. P., Crosby, E. J. & Hartman, Z. C. Cancer vaccine strategies using self-replicating RNA viral platforms. Cancer Gene Ther. 30, 794–802 (2023).

26. Lundstrom, K. Application of DNA replicons in gene therapy and vaccine development. Pharmaceutics 15, 947 (2023).

27. Blakney, A. K., Ip, S. & Geall, A. J. An update on self-amplifying mRNA vaccine development. Vaccines 9, 97 (2021).

28. Bloom, K., Berg, F. van den & Arbuthnot, P. Self-amplifying RNA vaccines for infectious diseases. Gene Ther. 28, 117–129 (2021).

29. Schmidt, C. & Schnierle, B. S. Self-amplifying RNA vaccine candidates: alternative platforms for mRNA vaccine development. Pathogens 12, 138 (2023).

30. Pijlman, G. P., Suhrbier, A. & Khromykh, A. A. Kunjin virus replicons: an RNA-based, non-cytopathic viral vector system for protein production, vaccine and gene therapy applications. Expert Opin. Biol. Ther. 6, 135–145 (2006).

31. Rappaport, A. R. et al. A shared neoantigen vaccine combined with immune checkpoint blockade for advanced metastatic solid tumors: phase 1 trial interim results. Nat. Med. 30, 1013–1022 (2024).

32. Komdeur, F. L. et al. First-in-human phase I clinical trial of an SFV-based RNA replicon cancer vaccine against HPV-induced cancers. Mol. Ther. 29, 611–625 (2021).

33. Morse, M. A. et al. Clinical trials of self-replicating RNA-based cancer vaccines. Cancer Gene Ther. 30, 803– 811 (2023).

34. Luke, J. J. et al. A phase I trial of intratumoral STX-001: A novel self-replicating mRNA expressing IL-12 alone or with pembrolizumab in advanced solid tumors. Journal of Clinical Oncology 42, TPS2696–TPS2696 (2024).

35. Hồ, N. T., et al. Safety, immunogenicity and efficacy of the self-amplifying mRNA ARCT-154 COVID-19 vaccine: pooled phase 1, 2, 3a and 3b randomized, controlled trials. Nat. Commun. 15, 4081 (2024).

36. Saraf, A. et al. An Omicron-specific, self-amplifying mRNA booster vaccine for COVID-19: a phase 2/3 randomized trial. Nat. Med. 30, 1363–1372 (2024).

37. Oda, Y. et al. Immunogenicity and safety of a booster dose of a self-amplifying RNA COVID-19 vaccine (ARCT-154) versus BNT162b2 mRNA COVID-19 vaccine: a double-blind, multicentre, randomised, controlled, phase 3, non-inferiority trial. Lancet Infect. Dis. 24, 351–360 (2024).

38. Cafferty, S. M. et al. In vivo validation of a reversible small molecule-based switch for synthetic self-amplifying mRNA regulation. Mol. Ther. 29, 1164–1173 (2021).

39. Ramadurgum, P. & Hulleman, J. D. Protocol for designing small-molecule-regulated destabilizing domains for in vitro use. STAR Protoc. 1, 100069 (2020).

40. Banaszynski, L. A., Chen, L., Maynard-Smith, L. A., Ooi, A. G. L. & Wandless, T. J. A rapid, reversible, and tunable method to regulate protein function in living cells using synthetic small molecules. Cell 126, 995–1004 (2006).

41. Abraham, R., et al. ADP-ribosyl–binding and hydrolase activities of the alphavirus nsP3 macrodomain are critical for initiation of virus replication. Proc. Natl. Acad. Sci. 115, E10457–E10466 (2018).

42. Hyde, J. L. et al. The 5′ and 3′ ends of alphavirus RNAs – Non-coding is not non-functional. Virus Res. 206, 99–107 (2015).

43. Rupp, J. C., Sokoloski, K. J., Gebhart, N. N. & Hardy, R. W. Alphavirus RNA synthesis and non-structural protein functions. J. Gen. Virol. 96, 2483–2500 (2015).

44. Shin, G. et al. Structural and functional insights into alphavirus polyprotein processing and pathogenesis. Proc. Natl. Acad. Sci. 109, 16534–16539 (2012).

45. Law, Y.-S. et al. Interdomain Flexibility of Chikungunya Virus nsP2 Helicase-Protease Differentially Influences Viral RNA Replication and Infectivity. J. Virol. 95, 10.1128/jvi.01470-20 (2020).

46. Yıldız, A., Răileanu, C. & Beissert, T. Trans-amplifying RNA: a journey from alphavirus research to future vaccines. Viruses 16, 503 (2024).

47. Götte, B., Liu, L. & McInerney, G. M. The enigmatic alphavirus non-structural Protein 3 (nsP3) revealing its secrets at last. Viruses 10, 105 (2018).

48. Vasiljeva, L. et al. Regulation of the sequential processing of Semliki Forest virus replicase polyprotein. J. Biol. Chem. 278, 41636–41645 (2003).

49. Shirako, Y. & Strauss, J. H. Regulation of Sindbis virus RNA replication: uncleaved P123 and nsP4 function in minus-strand RNA synthesis, whereas cleaved products from P123 are required for efficient plus-strand RNA synthesis. J. Virol. 68, 1874–1885 (1994).

50. DuPage, M., Dooley, A. L. & Jacks, T. Conditional mouse lung cancer models using adenoviral or lentiviral delivery of Cre recombinase. Nat. Protoc. 4, 1064–1072 (2009).

51. Wang, J., Isaacson, S. A. & Belta, C. Modeling genetic circuit behavior in transiently transfected mammalian cells. ACS Synth. Biol. 8, 697–707 (2019).

52. Wang, J., Isaacson, S. A. & Belta, C. Predictions of genetic circuit behaviors based on modular composition in transiently transfected mammalian cells. 2018 IEEE Life Sci. Conf. (LSC) **00**, 85–88 (2018).

53. Davidsohn, N. et al. Accurate predictions of genetic circuit behavior from part characterization and modular composition. ACS Synth. Biol. 4, 673–681 (2015).

54. Lautaoja, J. H. et al. Differentiation of murine C2C12 myoblasts strongly reduces the effects of myostatin on intracellular signaling. Biomolecules 10, 695 (2020).

55. Kislinger, T. et al. Proteome Dynamics during C2C12 Myoblast Differentiation* S. Mol. Cell. Proteom. 4, 887– 901 (2005).

56. Broderick, K. E. & Humeau, L. M. Enhanced Delivery of DNA or RNA Vaccines by Electroporation. in 193–200 (Springer New York, New York, NY, 2017). doi:10.1007/978-1-4939-6481-9_12.

57. Chaudhary, N., Weissman, D. & Whitehead, K. A. mRNA vaccines for infectious diseases: principles, delivery and clinical translation. Nat. Rev. Drug Discov. 20, 817–838 (2021).

58. Blau, H. M. & Springer, M. L. Muscle-mediated gene therapy. 333, 1554–1556.

59. Lu, Q. L., Bou-Gharios, G. & Partridge, T. A. Non-viral gene delivery in skeletal muscle: a protein factory. Gene Ther. 10, 131–142 (2003).

60. Bhagchandani, S. H. et al. Two-dose priming immunization amplifies humoral immunity by synchronizing vaccine delivery with the germinal center response. Science Immunology 9, eadl3755 (2024).

61. Allen, C. D. C., Okada, T. & Cyster, J. G. Germinal-center organization and cellular dynamics. Immunity 27, 190–202 (2007).

62. Victora, G. D. & Nussenzweig, M. C. Germinal centers. Annu. Rev. Immunol. 429–57 (2012).

63. Gitlin, A. D., Shulman, Z. & Nussenzweig, M. C. Clonal selection in the germinal centre by regulated proliferation and hypermutation. Nature 509, 637–640 (2014).

64. Jones, R., Bragagnolo, G., Arranz, R. & Reguera, J. Capping pores of alphavirus nsP1 gate membranous viral replication factories. Nature 589, 615–619 (2021).

65. Jones, R., et al. Structural basis and dynamics of Chikungunya alphavirus RNA capping by nsP1 capping pores. Proc. Natl. Acad. Sci. 120, e2213934120 (2023).

66. Bakar, F. A. & Ng, L. F. P. Nonstructural proteins of alphavirus—potential targets for drug development. Viruses 10, 71 (2018).

67. Kril, V. et al. Alphavirus nsP3 organizes into tubular scaffolds essential for infection and the cytoplasmic granule architecture. Nat. Commun. 15, 8106 (2024).

68. Pietilä, M. K., Hellström, K. & Ahola, T. Alphavirus polymerase and RNA replication. Viral polymerases 234, 44–57 (2017).

69. Tan, Y. B. et al. Molecular architecture of the Chikungunya virus replication complex. Sci. Adv. 8, eadd2536 (2022).

70. Peng, H. et al. Non-antibiotic small-molecule regulation of DHFR-based destabilizing domains in vivo. Mol. Ther. - Methods Clin. Dev. 15, 27–39 (2019).

71. Wroblewska, L. et al. Mammalian synthetic circuits with RNA binding proteins for RNA-only delivery. 33, 839– 841 (2015).

72. Li, B. et al. An orthogonal array optimization of lipid-like nanoparticles for mRNA delivery in vivo. Nano Lett. 15, 8099–8107 (2015).

73. An, H., Landis, J., T., Bailey, A., G., Marron, J., S. & Dittmer, D., P. dr4pl: a stable convergence algorithm for the 4 parameter logistic model. R J. 11, 171 (2019).

