## Supplementary Information for "Engineering gene expression dynamics via self-amplifying RNA with drug-responsive non-structural proteins"

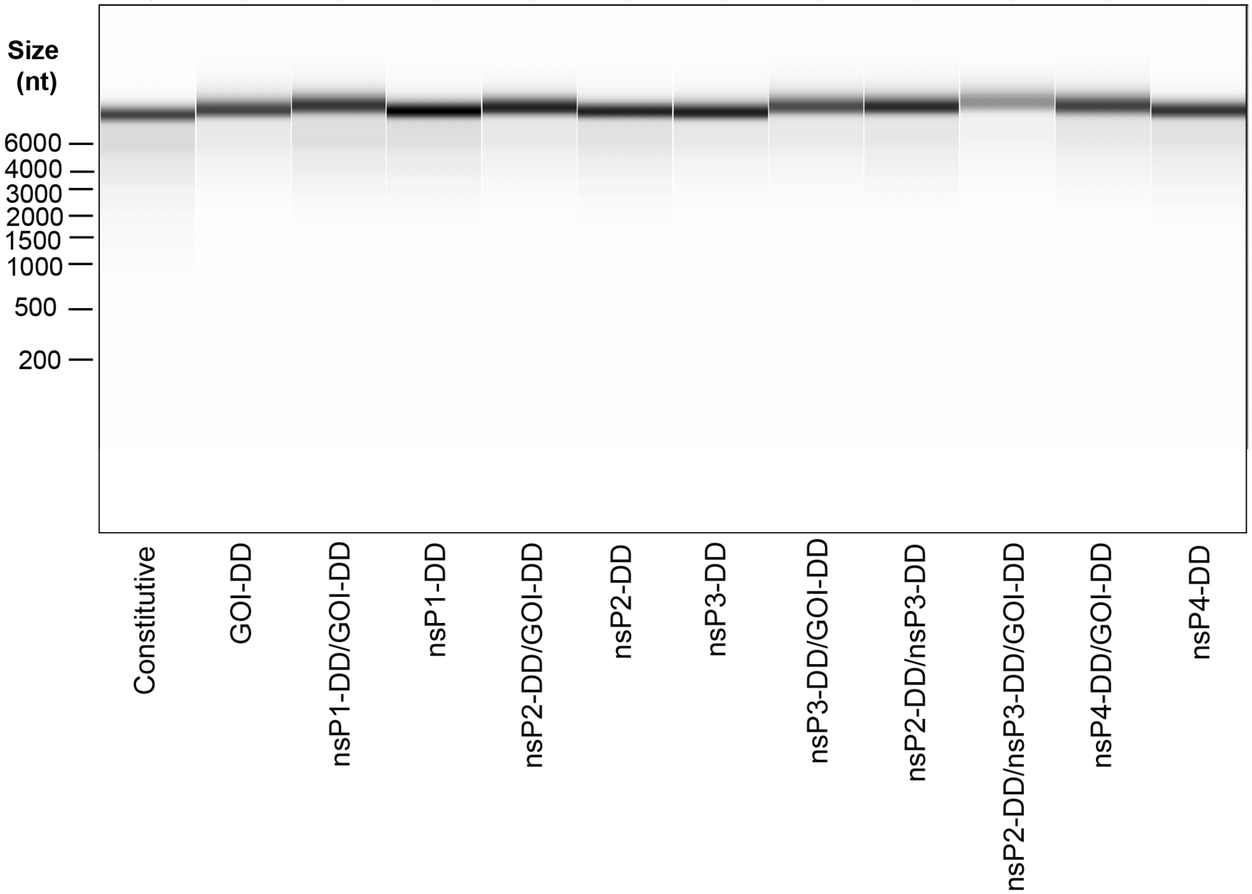


**Supplementary Figure S1. Quality of saRNAs synthesized by in vitro transcription.** saRNA samples with different fLuc-expressing gene circuits were analyzed on a Bioanalyzer. Shown is the resulting virtual gel image, which demonstrates a clear, single band for each sample, indicating high purity and intact RNA without significant degradation.

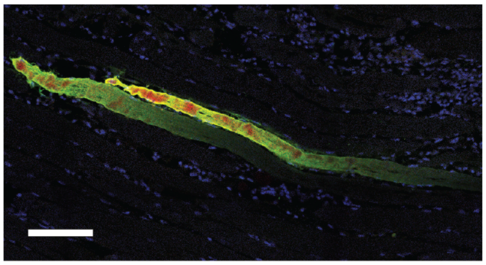

**Supplementary Figure S2: Transgene expression from saRNAs in mouse muscles.** Confocal microscopy images of muscle cryosections**.** Gastrocnemius muscles were harvested from mice three days post intramuscular injection with saRNAs encoding transmembrane mCherry. The muscles were flash-frozen in OCT compound, cryosectioned, and stained with DAPI for nuclear staining and an anti-mCherry antibody conjugated to Alexa488. The confocal microscopy image shows the localization of mCherry expression (red) and Alexa488 (green) as well as nuclei (blue) within the muscle tissue. Scale bar 100 µm.

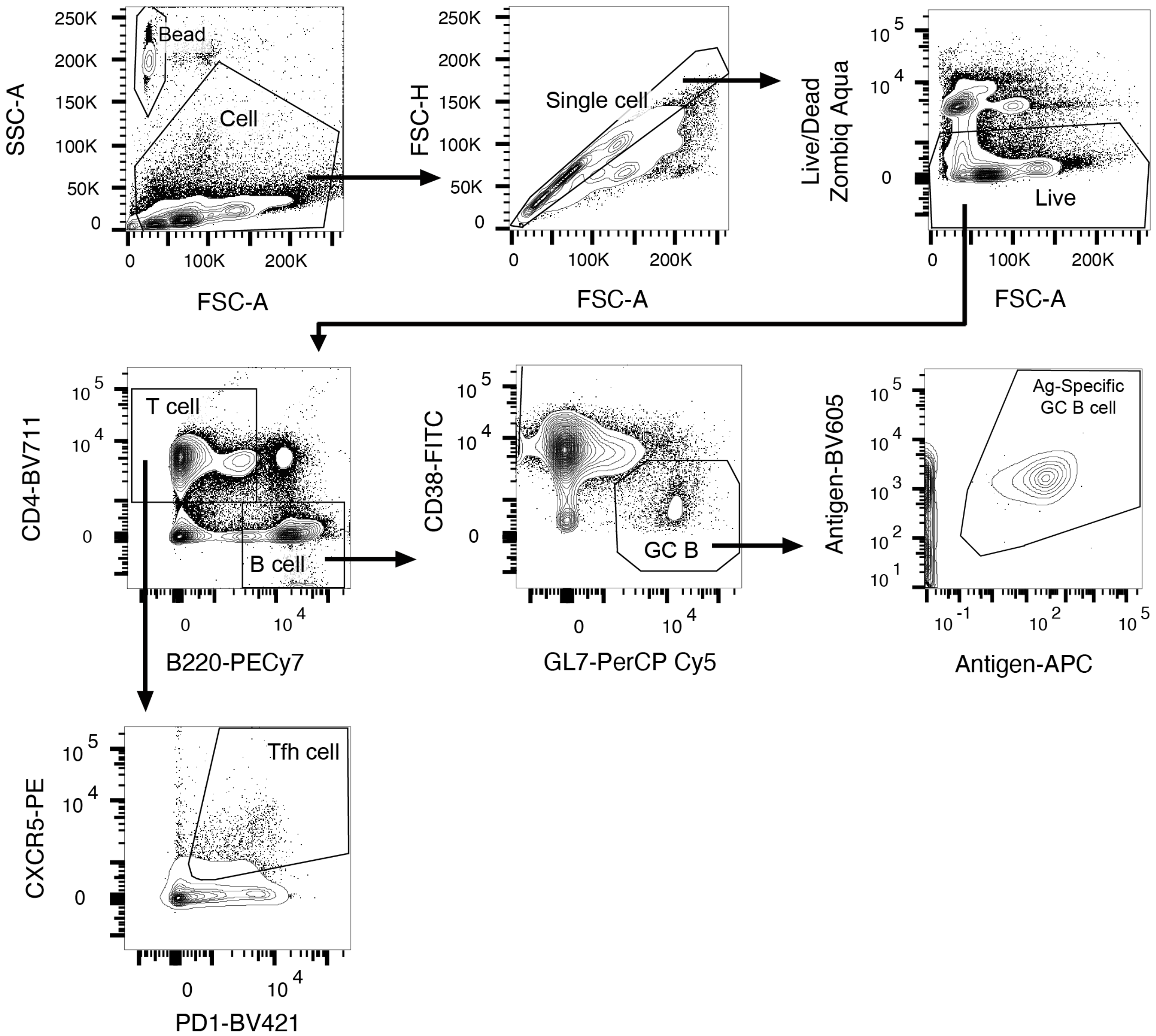


**Supplementary Figure S3: Flow cytometry gating strategy for GC Tfh and antigen-specific B cells analysis.** Lymphocytes isolated from lymph nodes were stained with the indicated markers. Live cells (Zombie Aqua negative) were pre-gated on FSC-A/FSC-H to select singlets and then gated on CD4 and B220 markers. B cells (CD4- B220+) were gated for low CD38 and high GL7 expression to identify GC B cells, followed by gating with MD39-tetramers to select for antigen specificity. T cells (CD4+ B220-) were gated for high PD1 and CXCR5 expression to identify Tfh cells.

**Supplementary Table 1: Kinetic parameters of the GOI-DD and nsP-DD circuits.**

| Circuit | $k_0$ | $k_{max}$ | $EC_{50}$ | $H$ |
|---|---|---|---|---|
| **GOI-DD** | 25.99 | 2151.12 | 35.77 | 1.70 |
| **nsP1-DD** | 1.75 | 359.41 | 3715.08 | 0.92 |
| **nsP2-DD** | 368.82 | 2168.31 | 1.45 | 2.19 |
| **nsP3-DD** | 242.51 | 1635.02 | 19.82 | 0.78 |
| **nsP2-DD/nsP3-DD** | 2.71 | 2992.64 | 8.76 | 1.84 |
